# Coevolution with Spatially Structured Rice Landraces Maintains Multiple Generalist Lineages in the Rice Blast Pathogen

**DOI:** 10.1101/2021.12.15.472812

**Authors:** Sajid Ali, Pierre Gladieux, Sebastien Ravel, Henri Adreit, Isabelle Meusnier, Joelle Milazzo, Sandrine Cros-Arteil, François Bonnot, Baihui Jin, Thomas Dumartinet, Florian Charriat, Alexandre Lassagne, Xiahong He, Didier Tharreau, Huichuan Huang, Jean-Benoît Morel, Elisabeth Fournier

**Author notes:** Corresponding authors &.

## Abstract

Traditional agrosystems, where humans, crops and microbes have coevolved over long periods, can serve as models to understand the eco-evolutionary determinants of disease dynamics and help the engineering of durably resistant agrosystems. Here, we investigated the genetic and phenotypic relationship between rice (*Oryza sativa*) landraces and their rice blast pathogen (*Magnaporthe oryzae*) in the traditional Yuanyang terraces of flooded rice paddies in China, where rice landraces have been grown and bred over centuries without significant disease outbreaks. Analyses of genetic subdivision revealed that *indica* rice plants clustered according to landrace names. Three new diverse lineages of rice blast specific to the Yuanyang terraces coexisted with lineages previously detected at the worldwide scale. Population subdivision in the pathogen population did not mirror pattern of population subdivision in the host. Measuring the pathogenicity of rice blast isolates on landraces revealed generalist life histories. Our results suggest that the implementation of disease control strategies based on the emergence or maintenance of a generalist lifestyle in pathogens may sustainably reduce the burden of disease in crops.

## INTRODUCTION

A major challenge for modern agriculture is to implement sustainable solutions ensuring food security by promoting crop health while decreasing our reliance on agrochemicals (Tilman, 2011). Globalization and agricultural intensification have disrupted the coevolutionary battle in which plants and pathogens engage in natural ecosystems, generally favoring larger pathogen population sizes (i.e., more widespread and intense epidemics) and thus rapid evolution of pathogen aggressiveness and infectivity (Burdon and Thrall, 2008; Gladieux et al., 2015; Parker and Gilbert, 2004). Monocultures of varieties bred for high yield and disease resistance are also vulnerable to disease outbreaks, because they impose strong directional selection on pathogens, and because mutants that can overcome resistance in one individual plant can infect all plants in a field and hence quickly spread (Hill, 2001; Stukenbrock and McDonald, 2008; Zhan et al., 2015). The literature in plant pathology provides many examples of so-called boom-and-bust disease dynamics, in which newly deployed resistant varieties are rapidly colonized by pathogen variants able to overcome new resistance genes (Brown, 1994; de Vallavieille-Pope et al., 2012; Guérin et al., 2007). In contrast, in unmanaged, natural, ecosystems, pathogen prevalence is generally lower, and disease epidemics more limited in time and space (Burdon and Thrall 2014). Long-term empirical studies and modeling work suggest that ecological and environmental heterogeneity, with highly patchy and variably diverse host plant populations, varying abiotic conditions and the co-occurrence of closely related but distinct or phylogenetically-distant plants can contribute to limit the burden of disease in the wild (Burdon and Thrall, 2008; Zhan et al., 2015).

Metapopulation dynamics and frequency dependent selection in heterogeneous environment create a mosaic of local coevolutionary scenarios ranging from local adaptation to maladaptation (Laine, 2007), depending on the biology of the system.

Traditional agrosystems are promising models for deriving new disease management rules for modern agrosystems (Chentoufi et al., 2014; Sahri et al., 2014). Transfering knowledge gained from studies of the mechanisms underlying the stability of plant-pathogen associations in the wild (Burdon and Thrall, 2008) is hindered by divergence in the structure and complexity of unmanaged ecosystems and modern agrosystems caused by marked differences in the impact of humans on the spatio-temporal distribution of host diversity between the two types of systems. Unlike modern agrosystems and modern crops, which have been engineered and intensely selected to improve yield and quality under relatively low-stress conditions, landraces and their agrosystems have been selected and developed for their capacity to provide stable yields in specific environmental conditions and under low-input agriculture. The value of landraces as sources of genetic variation, or the value of traditional agrosystems as models for re-engineering modern agrosystems, are generally accepted (Feuillet et al., 2008). However, there has been remarkably little effort to investigate causal links between the structure of genetic and phenotypic diversity in crops and pathogens on the one hand and disease dynamics on the other.

The traditional, centuries-old agrosystem of the Yuanyang terraces (YYT) of flooded rice paddies (Yunnan, China) represents an outstanding model system to investigate the factors that render plant agrosystems less conducive to disease (Liao et al., 2016). More than 180 landraces, mostly *indica* rice, have been grown for centuries in the Yuanyang terraces (Gao et al., 2012; Jiao et al., 2012; Yang et al., 2017). The Yuanyang landraces are notorious for being little affected by diseases (Sheng, 1990), such as rice blast caused by *Pyricularia oryzae* (syn., *Magnaporthe oryzae*), which is the most important rice disease worldwide (Dean et al., 2012).

Rice blast is widely spread on all ecotypes of rice and in different ecological zones, where it has a massive socio-economic impact on human populations (Dean et al., 2012; Tharreau et al., 2009). Rice blast is caused by one out of several host-specific lineages of *P. oryzae* (Gladieux et al., 2018). The rice-specific lineage is subdivided in three clonal (Gladieux et al., 2018)and one recombining and genetically more diverse lineage mainly distributed in Southeast Asia (Gladieux et al., 2018; Saleh et al., 2014). Cross inoculation experiments with globally distributed isolates pathogenic on rice have revealed host specialization of *P. oryzae* to the main groups of modern rice varieties (Gallet *et al*., 2016; Gladieux *et al*., 2018b). In the traditional YYT agrosystem, local adaptation to *indica* and *japonica* host ecotypes was also observed, and was associated with major differences in basal and effector-triggered immunity in the host (Liao et al., 2016). However, the coevolutionary interactions underlying the overall lower disease burden observed in YYT, remains unknown.

In this study we addressed whether the lower disease pressure observed on *indica* landraces, which represent 90 % of acreage in YYT, could result from the elevated landrace diversity extant in YYT, mitigating the emergence of *P. oryzae* populations specialized to *indica* landraces. We first analysed the population structure of YYT rice landraces on the one hand and *P. oryzae* populations on the other hand. We then used paired samples of *P. oryzae* pathogens and their plants of origin to address whether host and pathogen populations were genetically co-structured in order to establish if *P. oryzae* genotypes were specialized to their native host genotypes.

## MATERIALS AND METHODS

### Selection of rice accessions

In September 2014 and 2015, we collected plants just before harvest in two villages in YYT i.e., Jingkou and Xiaoshuijing (Supplementary Information SI1 Table SI1.1) from the six most popular rice landraces in the area (20 diseased plants + 10 healthy plants/ field, two fields / variety in each village). Panicles with mature seeds were kept for all plants so that each plant accessions could be selfed and grown again for DNA extraction and/or for multiplication. From this sampling, a set of 92 rice accessions representing indiginous landraces (Table SI1.1), was genotyped using Genotyping-by-Sequencing (Arbealz et al., 2015). DNA for these 92 rice accessions was extracted from individual plants. Half a leaf of each plant was ground into powder in liquid nitrogen. A volume of 750 ul of pre-warmed extraction buffer (CTAB 2% w/v, Tris-HCl 200 mM pH 8.0, EDTA 20 mM pH 8.0, NaCl 1.4 M, Polyvinylpyrrolidone (K30) 1% w/v, β-mercaptoethanol 1% v/v) was added to the powder and incubated at 65°C for 45 min. After centrifugation 15 min at 13000 rpm, the supernatant was recovered and extracted with the same volume of dichloromethane: isoamyl-alcohol (24:1). After centrifugation 15 min at 13000 rpm, the resulting supernatant was treated with RNAse A (0,1 ug/ml final) for 20 min at 37°C. The remaining nucleic acids were precipitated with cold isopropanol for 20 min at −20°C and centrifuged 10 min at 15000 rpm. The DNA pellet was washed once with ethanol 76%:sodium acetate 200 mM, once with ethanol 76%: sodium acetate 10 mM and finally resuspended in TE (Tris-HCl 10mM, EDTA 1mM pH8) buffer. DNA quality and quantity were checked using Nano-drop, Qubit^®^ dsDNA BR Assay Kits and on agarose gel. Library preparation and sequencing were performed at UMR AGAP (Montpellier, France) following the description in (Elshire et al., 2011). DNA was digested with ApeKI for library preparation and subsequent sequencing with an Illumina Genome Analyzer II (San Diego, California, USA).

To evaluate the genetic distance between YYT landraces and worldwide representatives of various rice sub-species, we selected 113 accessions of the worldwide rice sequencing data studied in Huang *et al*. (2012) and 103 accessions from the study of Wang *et al*. (2017) from the European Nucleotide Archive (ENA) database, chosen to maximize the geographic origins, sub-species representatives and genetic diversity (Supplementary information SI1).

### Processing of genomic data and SNP calling for rice accessions

Demultiplexing of raw GBS data, mapping and SNP calling were implemented in a pipeline using Toggle v0.3.3 (Monat et al., 2015). Reads were demultiplexed with PROCESSRADTAGS and mapped to the IRGSP-1.0 Nipponbare reference genome (Kawahara et al. 2013) using BWA (Li and Durbin, 2009) with option –n 5 for sub-commands aln and SAMSE. The alignments were sorted with picardToolsSortSam and SamtoolsView (http://broadinstitute.github.io/picard/, Li 2011). The GATK suite (McKenna et al. 2017) was used for downstream treatments. We used Realignertargetcreator to define suitable intervals for local realignments and Indelrealigner to perform local realignment of reads around indels. Markduplicates was used to remove duplicates, available in Picardtools. The output bam files were divided into per chromosome bam files with Bamtools. SNP calling was made with GATK for each chromosome with Gatkhaplotypecaller, while filtering sites with the option Badcigar. High-confidence SNPs were identified using GATK’s Variantfiltration to filter variants based on parameters DP>10, QUAL > 30.

Genomic data from the 216 worldwide rice accessions were mapped against the IRGSP-1.0 Nipponbare reference genome using the same procedure. Mapping data were post-processed as described above. Analyses were conducted on the intersect between the set of SNPs identified with GBS data for YYT landraces and the one identified with whole genome data for worldwide accessions.

### Population genetic analysis of genomic variation in rice

In the total sample containing 92 YYT and 216 worldwide rice accessions, population subdivision was assessed without considering *a priori* information about the name, rice type or location of origin of rice samples with FastStructure, by varying the number of clusters (K) from 2 to 10 (Raj et al., 2014), using BED and BIM files generated with PLINK (Purcell et al., 2017). Genetic clustering was also inferred for the 46 rice accessions paired with corresponding *P. oryzae* isolates using sNMF analyses by generating GenLight object in R-statistical environment, and running the analyses by varying the number of clusters (K) from 2 to 10. We inferred the genealogical relationships between these 46 accessions with RAxML (Stamatakis 2014), based on pseudo-assembled genomic sequences (i.e. genomic sequences generated from the table of SNPs and reference sequences). We used the General Time-Reversible model of nucleotide substitution with the Γ model of rate heterogeneity, and performed 100 bootstrap replicates to estimate branch support.

### *Sampling and isolation of* P. oryzae

Rice blast samples were collected between 2009 and 2016 in eight villages from YYT, including the two villages where rice landraces were sampled (Supplementary Information SI2, Fig. SI2.1), just before harvest. Diseased organs (leaves or collars) were kept in paper bags and dried at room temperature. Genetically pure isolates of *P. oryzae* were obtained after single spore isolation from colonies grown from infected plant material placed in humid chamber at 21°C for 1–2 days. Single-spored fungal isolates were then grown on rice flour medium, as previously described (Silué and Nottéghem 1990), and stored on filter paper at −20°C, as described by Valent *et al*. (1986).

### *Selection of* P. oryzae *isolates and rice genotypes for downstream analyses*

To infer the population structure of *P. oryzae* in YYT and place the diversity observed in YYT in the global diversity of *P. oryzae* pathogens infecting rice, we analysed a set of 512 isolates sampled in YYT, along with 45 samples representing the main lineages observed at world scale (Saleh *et al*., 2014; Gladieux *et al*., 2018b), which were all genotyped using 13 microsatellites (Adreit et al., 2007). Part of the microsatellite data was available from previous studies in the worldwide (Saleh *et al*., 2014; 45 isolates) and YYT (Liao *et al*., 2016; 214 isolates) contexts. To further characterize the genetic co-structure between hosts and pathogens, we analysed a subset of paired samples collected in 2015 and composed of 46 plants representative of the five most popular *indica* landraces in YYT, and their corresponding 46 *P. oryzae* pathogens (i.e. one isolate per plant coming from rice blast lesions found on the plant).

### *Genomic DNA extraction for re-sequencing of* P. oryzae *isolates*

To extract genomic DNA matching the quality criteria for full genome resequencing, the 46 *P. oryzae* isolates selected above were first grown on rice flour solid medium for mycelium regeneration, then in liquid rice flour medium following Adreit *et al*. (2007). Genomic DNA extraction was carried out using 100 mg of fresh mycelium from liquid culture. Fresh mycelium dried on Miracloth paper was crushed in liquid nitrogen. Nucleic acids were subsequently extracted with a lysis buffer (2 % CTAB - 1.4 M NaCl - 0.1 M Tris-HCl pH 8 - 20 mM EDTA pH 8 added before use with 1 % final of Na_2_SO_3_), then purified with a chloroform:isoamyl alcohol 24:1 treatment, precipited overnight in isopropanol, and rinsed with 70% ethanol. The extracted nucleic acids were further treated with RNase A (0.2mg/mL final) to remove RNA and purified with another chloroform:isoamyl alcohol 24:1 treatment followed by an overnight ethanol precipitation. The concentration of extracted genomic DNA was assessed on Qubit® using the dsDNA HS Assay Kit. The purity of extracted DNA was checked by verifying that the 260/280 and 260/230 absorbance ratios measured with NanoDrop were between 1.8 and 2.0. We also ran 0.5 ot 1 μg DNA extracts on agarose gel to visually verify the absence of RNAs and degraded DNAs. Preparation of sequencing libraries and Illumina HiSeq 2500 sequencing was performed at GenWiz Inc. USA, resulting in paired-end reads of 150 nucleotides with ca. 500 bp insert size.

### *Processing of genomic data and SNP calling for* P. oryzae *isolates*

As for rice genomic data, we used Toggle to implement a pipeline for raw reads processing, mapping and SNP calling. Raw reads were trimmed to remove barcodes, adapters and ambiguous base calls. Trimmed reads were mapped against reference genome 70-15 version 8 (Dean et al., 2005) using BWA with option -n 5 for sub-command aln and option -a 500 for paired-end analyses sub-command *sampe*. The alignments were sorted with PICARDTOOLSSORTSAM and SAMTOOLSVIEW (http://broadinstitute.github.io/picard/, Li 2011). Intervals to target for local realignment were defined using Realignertargetcreator, and local realignment of reads around indels were performed with Indelrealigner. Duplicates were removed with Markduplicates. SNPs were then called using the UnifiedGenotyper tool in GATK, while keeping all sites of the reference genome using the option Emit_all_sites. High-confidence SNPs were identified using GATK’s variantfiltration option with the following parameters: MQ0< 3.0 (total mapping quality zero reads), depth ≥ 15.0 (number of reference alleles + number of alternative alleles, computed as the sum of allelic depths for the reference and alternative alleles in the order listed), and RA ≤ 0.1 (number of reference alleles / number of alternative alleles).

### *Population genetic analysis in* P. oryzae

To detect genetic lineages within the 46 *P. oryzae* isolates for which we had full-genome information, we combined these data with 48 worldwide rice-infecting *P. oryzae* genomes published by Gladieux *et al*. (2018b). In Gladieux’s study, six rice-infecting lineages were described, two of which (lineages 5 and 6) being represented by one isolate each. Since then, two studies based on larger sets of fully sequenced and/or genome-wide genotyped isolates (Latorre et al., 2020; Thierry et al., 2021) showed that lineages 5 and 6 are in fact part of lineage 1. We used the phylogenetic network approach neighbor-net as implemented in Splitstree 4.13 (Bryant and Moulton, 2004). This allowed visualizing evolutionary relationships, while taking into account the possibility of recombination within or between lineages. We also assessed the genealogical relationships among the 46 YYT fully-sequenced *P. oryzae* isolates by analyzing pseudo-assembled genomic sequences (i.e. genomic sequences generated from the table of SNPs and reference sequences) with RAxML (Stamatakis, 2014). We used the General Time-Reversible model of nucleotide substitution with the Γ model of rate heterogeneity, and performed 100 bootstrap replicates to assess branch support. To assess the population subdivision among the genomic data of 46 *P. oryzae* isolates, sNMF analyses were performed by generating GenLight object in R-statistical environment. To assess population subdivision without considering *a priori* information about the origin of samples, and without assuming random-mating, discriminant analyses of principal components (DAPC) were conducted on the microsatellite data for the 512 YYT and 45 worldwide isolates. DAPC analyses were done using the Adegenet package in R (Jombart et al., 2010), by varying the number of inferred genetic clusters (K) from 2 to 10.

For microsatellite data, a distance-based neighbour-joining tree was generated with Population (Langella, 2008), and within-population diversity and linkage equilibrium parameters were estimated using Poppr package in R-environment (Kamvar et al., 2014). Nucleotidic diversity (π) within lineages identified using full-genome resequencing data was estimated using the package Egglib 3.0.0b10 (De Mita and Siol, 2012) and divergence among lineages was estimated in 10kb windows using the dxy statistic as implemented in the scikit-allel (https://zenodo.org/record/4759368#.YW1duxBBzwQ). LD decay along the genome was assessed within each genetic lineage using PopLDdecay (Zhang et al., 2019). We determined the mating type of each resequenced isolate using a BLAST search of Mat1 and Mat2 idiomorphs sequences within each genome assembled de novo using ABySS 2.0 with default parameters (Jackman et al., 2017).

### Phenotyping of host-pathogen biological interactions

Cross-compatibility among rice / *P. oryzae* paired samples was assessed through cross inoculation experiments. One rice accession (HO-Q-F16) could not be included since seeds were lacking, and the corresponding *P. oryzae* isolate (CH1866) was also excluded. The remaining 45 rice accessions were separated in two trays, each tray containing 22 or 23 accessions plus the two highly susceptible rice accessions CO39 and Maratelli (Gallet et al. 2016) used as positive controls, with six rice seeds sown per variety (i.e. 144 or 150 plants per tray). As many batches as *P. oryzae* isolates (i.e. 45) of such two trays were prepared (i.e. 90 trays in total). Trays were inoculated four weeks after sowing when plants had 4-5 leaves, each batch of two trays was inoculated with a single *P. oryzae* isolate. The fungal inocula were composed of conidia suspensions at 50,000 spores/ml and 25,000 spores/ml for the first and second repetition of the experiment, respectively. Spore suspensions were supplemented with 0.5% gelatin (Gallet et al., 2016). An inoculation corresponded to the spraying of the spore suspension of one particular isolate on one batch of two trays. All inoculations were performed at the same date. Seven days after inoculation, symptoms were read on four plants for each blast genotype × rice genotype interaction; the four corresponding leaves were glued on sticky papers and scanned for subsequent scoring of symptoms. The experimental design was repeated twice. Twelve other rice accessions were removed from the analysis because they were difficult to multiply, did not grow well in our controlled conditions, could not be assigned to any rice genetic cluster (BJ-Q-B06, Fig. 3 left panel), or were of modern origin (HongYang accessions). We thus ended with a matrix of 33*33 rice / *P. oryzae* paired samples with complete results.

Qualitative interactions were noted on each leaf qualitative scale of 1-6: scores 1-2 corresponding to incompatible reactions, scores 3-6 corresponding to compatible reactions (Gallet et al., 2014). The percentage of compatible / incompatible reactions for each interaction was estimated by counting the total number of compatible / incompatible leaves among the total number of leaves scored over the two experimental repetitions. When less than four leaves were available in total, the data was considered as missing. We verified that the qualitative scores of the two independent replications were positively correlated (Supplementary Information SI3, Fig. SI3.2). To obtain quantitative measure of host-pathogen compatibility phenotype, the scanned images were analysed with Ebimage package implemented in R statistical environment (Pau et al., 2010). In-house scripts were used for calibration and image analyses (https://github.com/sravel/LeAFtool). Briefly, calibrations were made according to discriminant analysis of RGB composition of pixels chosen and classified by the user as lesion, leaf and background, and the resulting discriminant functions were used to assign pixels of the entire image to these three categories. Statistical analyses were performed using the nlme package in R environment. The studied variable was the percentage of diseased leaf area. After log-transformation of this variable (y = log(x + 0.15)), we performed a two-step analysis. First we performed an ANOVA considering only the positive controls to evaluate the respective effects of the following factors: “repetition”, “tray”, *“P. oryzae* isolate”, “rice accession”, and the interaction between the last two. We obtained significant effects for fungal isolate (F = 5.13, P = 2.4e^-10^, df = 32) and rice landrace (F = 416.9, P < 10^-16^, df = 1); the effect of tray was significant (F = 1.7, P = 0.003, df = 98) but neglectable compared to the effect of experimental replicate (F = 181.7, P < 10^-16^, df = 1), and was therefore ignored in subsequent data analyses. We then analysed the log-transformed variable for the rest of the matrix (excluding the positive controls) using an ANOVA considering two factors: “repetition” and “combination” (corresponding to each *P. oryzae* isolate × rice accession combination). Heatmaps of the adjusted value of the log-transformed variable were drawn using ggplot2 package in R environment.

## RESULTS

### Genetic structure and diversity in YYT rice accessions

The SNP dataset combining GBS data from 92 YYT accessions and full-genome data from 216 worldwide accessions (Supplementary Information SI1) contained 44,855 SNPs having less than 30% missing calls. Analysis of population subdivision using FastStructure (Fig. 1) showed that YYT rice accessions formed two main groups that differentiate at K=3, one closely related to the *indica* worldwide ecotype, the other one related to the *japonica* worldwide ecotype (Fig. 1). This was confirmed by the genealogical relationships among accessions inferred using RAxML (Fig. SI1.1). The FastStructure analysis also showed that from K=7 YYT genotypes clustered according to landrace names used by farmers, indicating that these landraces are ‘population’ varieties composed of distinct, closely related genotypes. Altogether, these analyses showed that, for both *indica* and *japonica* types, the YYT landraces formed a genetic pool that is related to, but distant from, the worldwide representatives.

**Figure 1.**
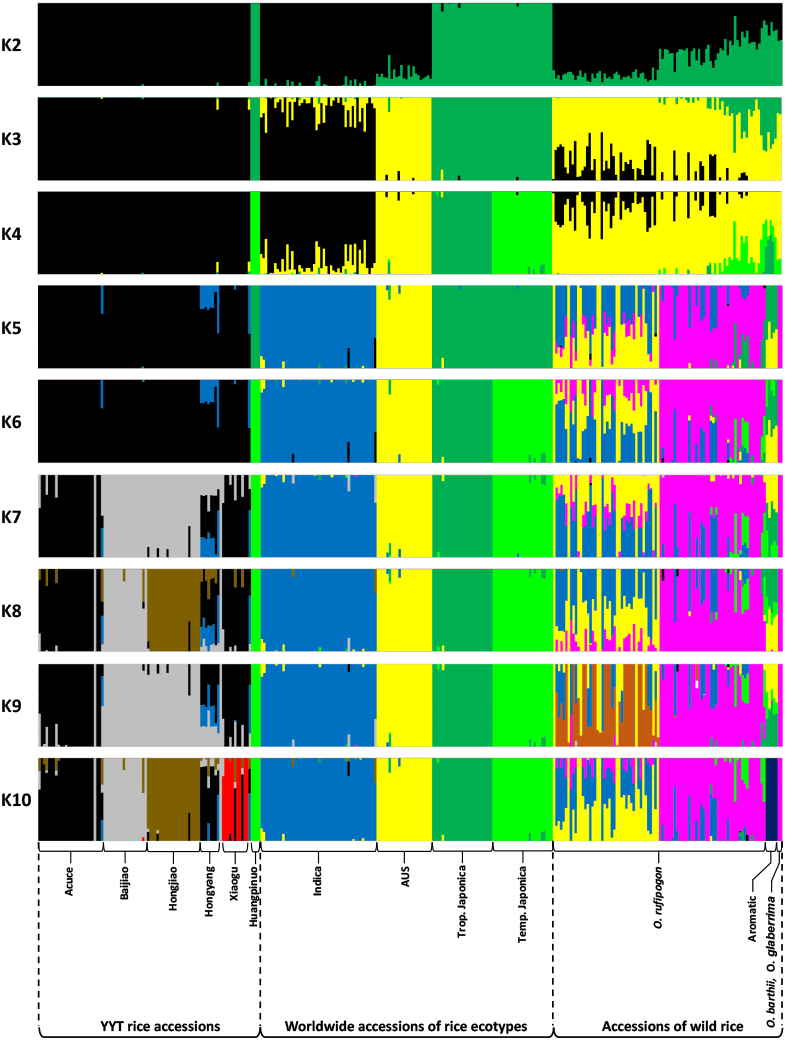
Population subdivision assessed with FastStructure within the total sample containing 92 YYT and 216 worldwide rice accessions. Each horizontal barplot represents subdivision assessed for one value of K, the number of inferred clusters, with K varying from 2 to 10. Vertical lines within each horizontal barplot represent the probability of ancestry of each individual in each inferred cluster. Vernacular names of YYT landraces and names of *O. sativa* subspecies and of wild *Oryza* species are indicated below the barplots.

### *Genetic diversity and recombination in* P. oryzae *lineages from YYT*

Analysis of population subdivision in *P. oryzae* revealed the coexistence in YYT of previously described worldwide lineages along with newly detected lineages specific to YYT. Both the NJ tree (Fig. 2A) and the DAPC barplot (Fig. 2B) estimated from microsatellite data for 513 YYT isolates and 44 worldwide representative isolates, showed that some YYT isolates grouped with previously described worldwide lineages (named W-lineages on the figures) whereas a large group of YYT isolates formed another group specific to this region, subdivided into several clusters. As expected from Liao et al. (2016), isolates coming from glutinous *japonica* landraces from YYT formed a separate clade, with some spillover genotypes on *indica* landraces (Fig. 2B). The coexistence of multiple *P. oryzae* lineages in YYT was further confirmed by whole-genome analysis of the 46 YYT isolates. Short-reads whole genome re-sequencing of these 46 isolates yielded an average coverage depth of 5X, that resulted in a final dataset of 66,102 SNPs without any missing data after mapping against the 70-15v8 reference genome. These data were pooled together with whole-genome SNPs data from 48 worldwide representative isolates previously sequenced (Gladieux et al., 2018) to build a neighbour-net network using Splitstree. This analysis confirmed that nine isolates from YYT were assigned to two of the four worldwide lineages previously described (Gladieux et al., 2018; Latorre et al., 2020; Thierry et al., 2021) (Fig. 2C: four isolates from YYT assigned to W-Lineage 1, five isolates from YYT assigned to W-Lineage 3), whereas the 37 remaining isolates were assigned to three well-supported lineages specific to YYT, hereafter named lineages YYT1 to 3 (Fig. 2C; Supplementary Information SI2 Table SI2.1). Our data thus supported a population subdivision into five genetic lineages.

**Figure 2.**
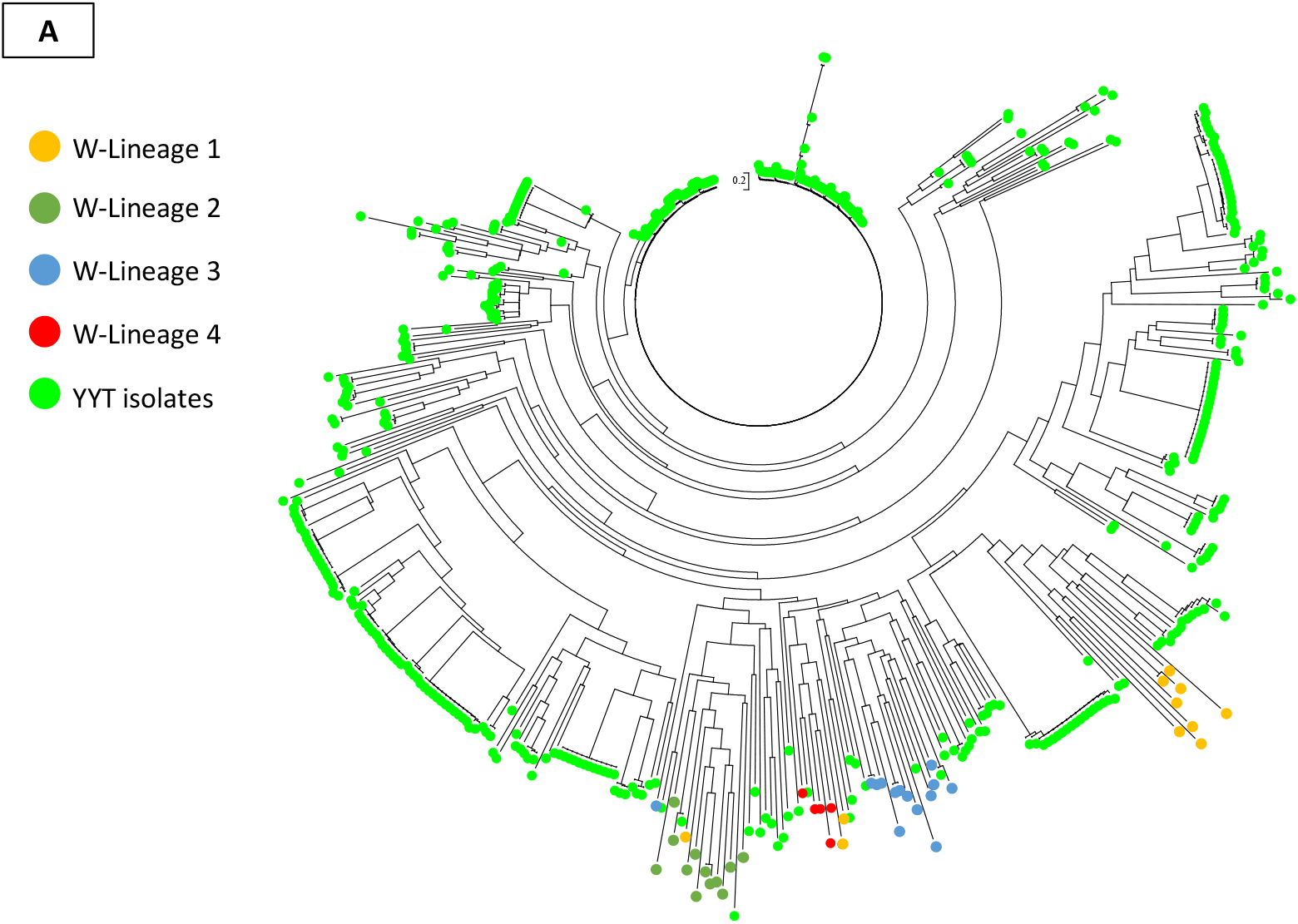
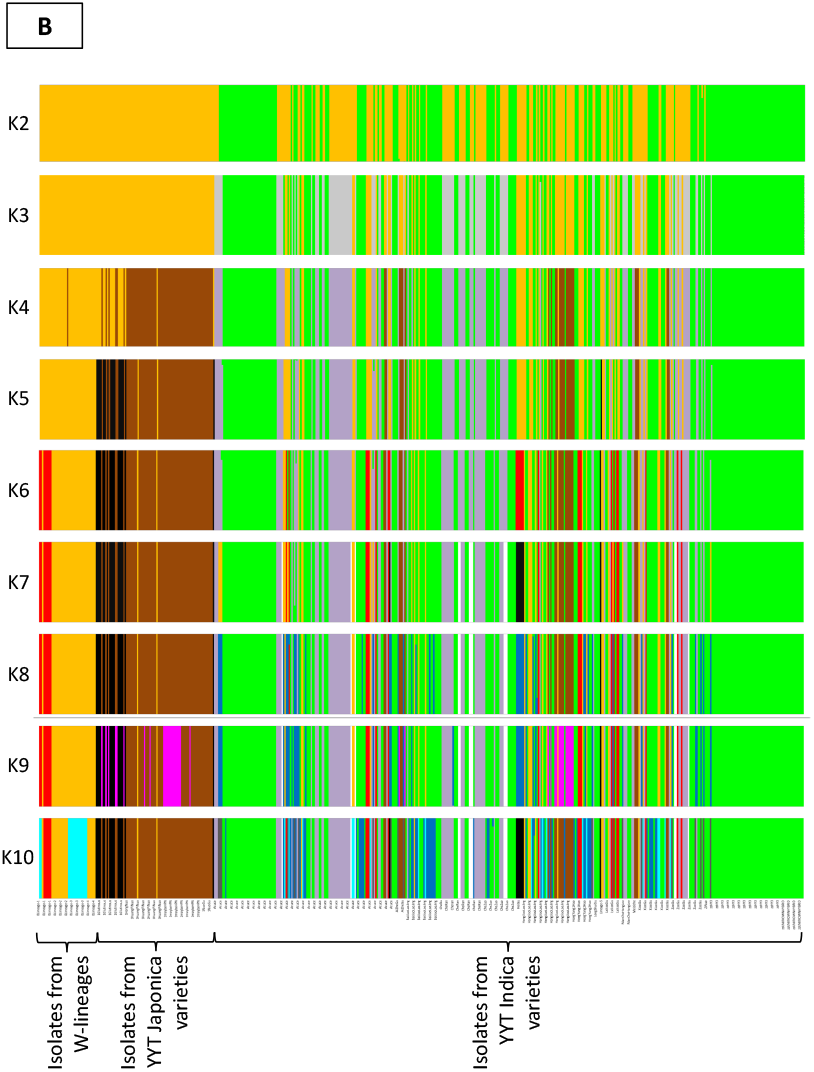
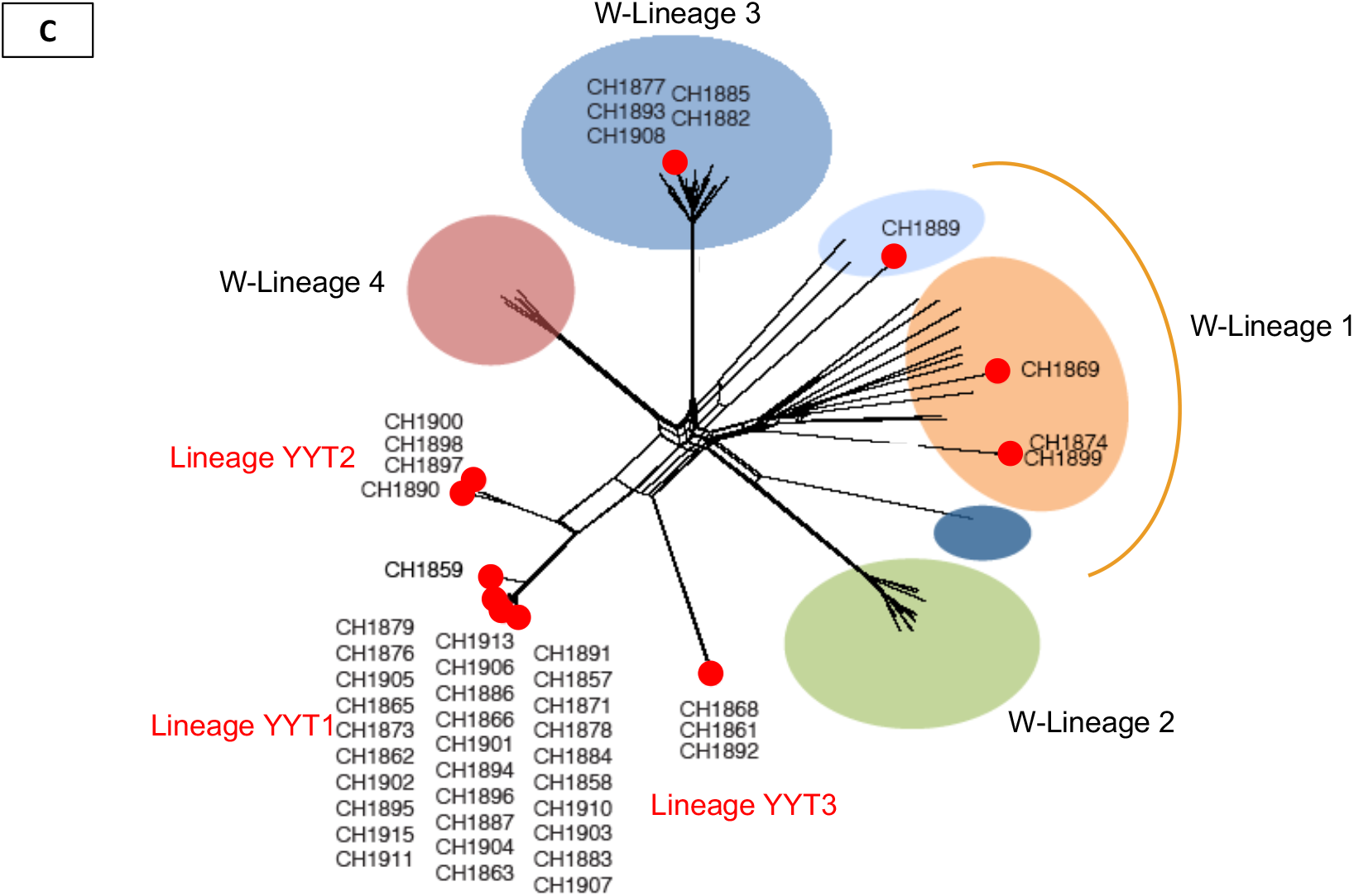
Analysis of subdivision of the *P. oryzae* population sampled in YYT. A. Phylogenetic relationships based on microsatellite data of 512 isolates from YYT (green dots) and 43 isolates representative of previously described worldwide lineages (W-Lineages, non-green dots). B. DAPC clustering analysis based on microsatellite data of 512 isolates from YYT and 43 isolates representative of previously described worldwide lineages. C. Phylogenetic network based on whole-genome SNPs data of 46 isolates from YYT (red dots) and 48 isolates representative of worldwide lineages (W-Lineages, non-dotted branches) inferred using the neighbor-net method. Worldwide isolates used in Fig. 2A, B and C were chosen to represent the four worldwide lineages defined in previous studies (Saleh *et al*., 2012; Gladieux *et al*., 2018b; Latorre *et al*., 2020; Thierry *et al*., 2021).

An overall high genetic variability was observed based on microsatellite data for the entire *P. oryzae* population sampled in YYT (513 isolates), as shown by gene diversity (0.566), Simpson’s diversity (0.947) and evenness (0.199). At the lineage level (Fig. 2C), nucleotide diversity estimated from full-genome SNPs data was the highest for YYT isolates from worldwide lineage 1 (π=2.00 × 10^-4^/bp) and was the lowest for isolates from lineage YYT3 (π=0.12 × 10^-4^/bp) (Table 1). As compared to π estimates in non-YYT isolates from worldwide lineages provided by Gladieux *et al*. (2018b), π estimates in YYT isolates assigned to worldwide lineages were either equal (for WL1: π=2.00 × 10^-4^/bp, this study, vs π=2.11 × 10^-4^/bp, Gladieux *et al.,* 2018b), or slightly higher (for WL3: π=0.53 × 10^-4^/bp, this study, vs π=0.45 × 10^-4^/bp, Gladieux *et al.,* 2018b). Sequence divergence among lineages was generally higher than nucleotide diversity within lineages, with dxy ranging between 0.09 × 10^-3^/bp between lineages YYT1 and YYT2 to 2.25 × 10^-3^/bp between lineages YYT2 and W-Lineage 3 (Table 1).

**Table 1.** Nucleotide diversity (π) within, and sequence divergence (dxy) between, genetic lineages estimated from whole-genome SNPs data for 46 *P. oryzae* isolates from YYT. Both statistics were estimated in non-overlapping 10kb windows and the average value per base pair is presented. W-lineages: worldwide lineages as described in Gladieux et al. (2018). Assignment of the 46 isolates from YYT to worldwide lineages is based on Fig. 2C & Table SI2.1.

| Lineage | Nucleotide diversity $\pi$ | Sequence divergence (dxy) | | | |
| --- | --- | --- | --- | --- | --- |
|  |  | W-lineage-3 | YYT1 | YYT2 | YYT3 |
| W-lineage-1 | $2.00 \times 10^{-04}$ | $2.27 \times 10^{-03}$ | $2.21 \times 10^{-03}$ | $2.20 \times 10^{-03}$ | $2.21 \times 10^{-03}$ |
| W-lineage-3 | $0.53 \times 10^{-04}$ | - | $2.23 \times 10^{-03}$ | $2.25 \times 10^{-03}$ | $2.14 \times 10^{-03}$ |
| YYT1 | $0.12 \times 10^{-04}$ | - | - | $0.09 \times 10^{-03}$ | $1.50 \times 10^{-03}$ |
| YYT2 | $0.22 \times 10^{-04}$ | - | - | - | $1.59 \times 10^{-03}$ |
| YYT3 | $0.12 \times 10^{-04}$ | - | - | - | - |

Signatures of recombination were searched within the *P. oryzae* population sampled in YYT. For the whole *P. oryzae* population sampled in YYT (513 isolates), the global linkage disequilibrium (r_D_) estimated from microsatellite data was 0.145: although significantly different from 0 (P-value=0.001; r_D_=0 is the expected value under the null hypothesis of free recombination), this value was far from 1 (expected value under complete clonality). The rate of sexual reproduction estimated from microsatellite data among these 513 isolates varied from 15% to 100% depending on the time period considered (Supplementary Information SI2 Table SI2.2)., Phi-tests performed within each lineage using whole-genome data were all significant, allowing to reject the null hypothesis of strict clonality. However, patterns of LD decay along the genome, estimated within each of the five lineages inferred in YYT, showed no evidence of recombination, although reticulations were observed in the minimum networks for lineage YYT1 and, to a lesser extent, for lineages WL1 and WL3 (Supplementary Information SI2, Fig. SI2.2). Finally, we searched the mating type genes in the *de novo* assembled genomes of the 46 fully sequenced isolates from YYT using BLAST, and confirmed the presence of both mating types in the whole *P. oryzae* sample. However, only isolates assigned to WL1 comprised both mating types, the other lineages carrying a single mating type (Supplementary Information SI2 Table SI2.3). Altogether, these results indicated that recombination events, if existing, did not leave significant signatures in this dataset.

### Lack of genetic co-structure between hosts and pathogens

To address the question of specialization to the host in *P. oryzae* population, we first compared the genetic structure of hosts and pathogens within the 46 paired samples of *indica* landraces and *P. oryzae* isolates from YYT. The total evidence tree built from rice GBS data (26,860 SNPs after removal of sites with missing; Fig. 3, left tree) showed that, as expected, YYT rice accessions clustered according to landrace vernacular name. The few exceptions to this clustering pattern could be explained by seed movements due to farmer practices at the time of harvest, resulting in variety mixture within paddies. Assignment analysis based on genomic data (Fig. 3, left barplots) further confirmed the clustering of YYT rice accessions by landrace name, and defined five main genetic clusters corresponding to Xiaogu, Hongyang, Acuce, Hongjiao and Baijao landraces. One accession (BJ_Q_B06) was considered as admixed. These results confirmed that landraces names in YYT correspond to well-defined genetic clusters, corresponding to landraces, i.e., populations of different - though genetically-related - genotypes.

**Figure 3.**
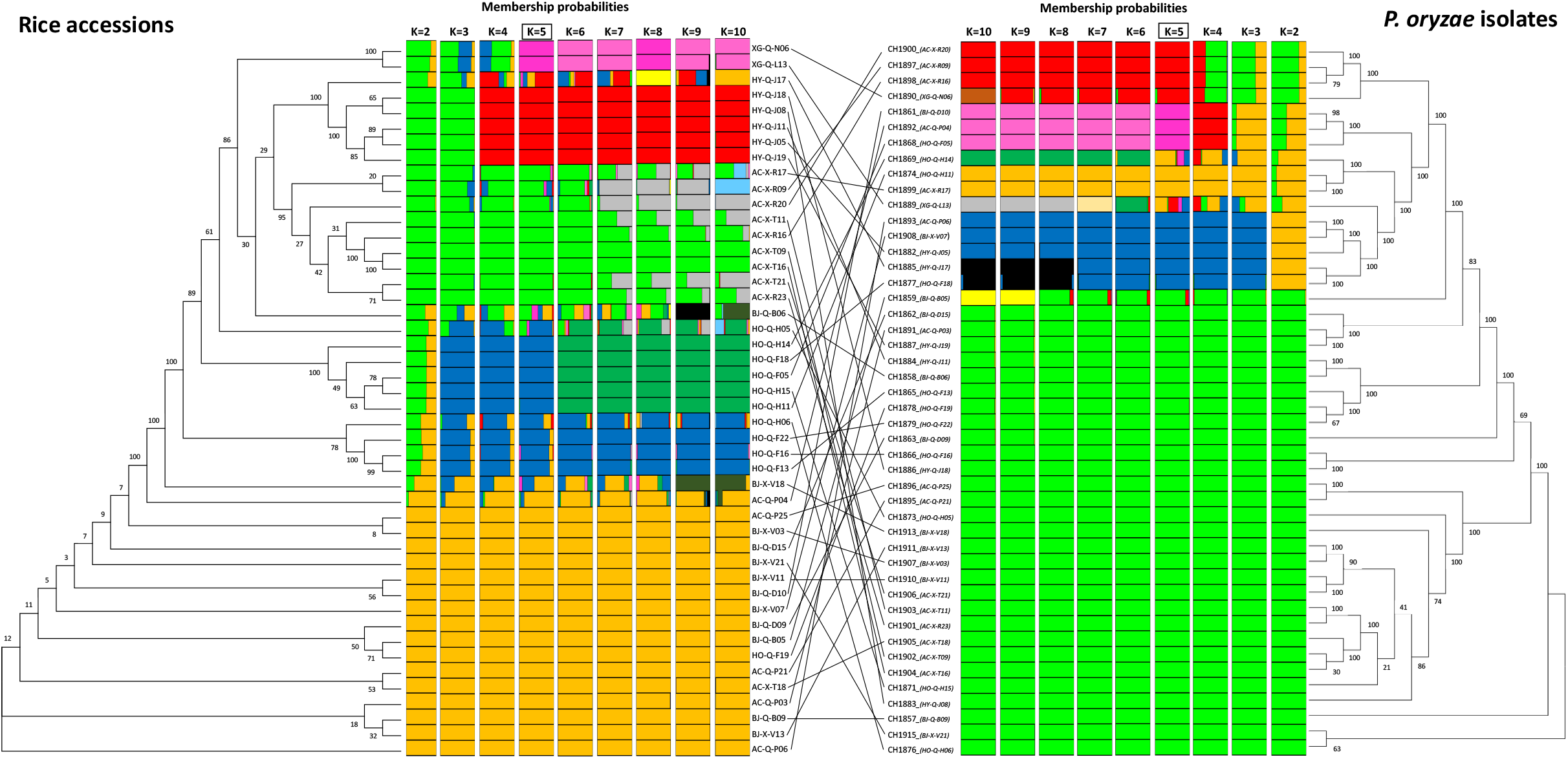
Comparison of genetic subdivision among 46 rice / *P. oryzae* paired samples. Left part of the figure: rice accessions analysed for 26,860 SNPs. Right part of the figure: corresponding *P. oryzae* isolates analysed for 66,102 SNPs. The genealogical trees were built using RAxML, with branch supports after 100 bootstrap replicates indicated on branches. The barplots show the result of DAPC clustering analysis for a number of genetic clusters (K) varying from 2 to 10 (each barplot indicates probabilities of ancestry of each individual in the corresponding K clusters). Black lines in the central part of the figure connect each rice accession to its corresponding paired *P. oryzae* isolate.

Population subdivision was also clearly evidenced for the pathogen, as shown both by the phylogenetic tree and the clustering analyses (Fig. 3, right panel). While considering five clusters (i.e. at K=5), the grouping of the 46 sequenced *P. oryzae* isolates was congruent with the subdivision previously inferred from the phylogenetic network (Fig. 2C): one group (yellow) encompassed three isolates assigned to worldwide lineages W-Lineage 1, one group (blue) encompassed five isolates assigned to the worldwide lineage 3, and the remining isolates were assigned to three groups specific to YYT (green, red and pink groups, corresponding respectively to lineages YYT1, 2 and 3 of Fig. 2C). Pairing host and pathogen samples in genome genealogies clearly showed a complete lack of genetic co-structure between host and pathogen populations (black lines in Fig. 3): pathogens did not clustered according to the landrace their host of origin belongs to, thus suggesting a lack of specialization to the host.

The lack of any subdivision explained by host in *P. oryzae* populations sampled on *indica* landraces from YYT was also confirmed by the DAPC analysis of microsatellite data, since genetic subdivision inferred from these data did not match the host of origin (Fig. 2B). Genetic diversity of pathogens estimated with microsatellite data was high on most rice landraces in YYT (except for HongYang 3 and an unknow landrace from which a high proportions of *P. oryzae* clones were sampled), ranging from 0.292 to 0.626 with a mean value of 0.445 (Supplementary Information 3, Table SI3.1), which compares to the ten most diverse populations described in Saleh *et al*. (2014). Among the 289 different microsatellite multilocus genotypes (MLGs) recovered from the 557 *P. oryzae* isolates sampled in YYT, 48 MLGs were detected more than once, 31 of which (64.5%) on multiple *indica* landraces (Table SI3.2). This confirmed that pathogen genotypes were distributed on all rice landraces.

### *Lack of specialization of* P. oryzae *populations to their hosts*

Cross-infection compatibility was tested for 33 rice plants and their paired *P. oryzae* isolates from YYT (1,089 possible combinations; Supplementary information SI3, Fig. SI3.1). Qualitative results showed a lack of phenotypic specificity for the vast majority of *P. oryzae* isolates to their native rice accession or to plants belonging to the same landrace as their native plant. Indeed, among the 1,082 combinations yielding analysable results (data were missing for 7 combinations), 1,025 interactions (94.7%) were fully compatible, only 3 (0.3%) were incompatible, and 54 (5%) were scored as undetermined. This result suggest that all major resistance genes present in this set of *indica* landraces genotypes are overcome by *P. oryzae*.

Analysis of quantitative interactions in the matrix, measured as the average diseased leaf area, revealed a lack of adaptation of *P. oryzae* to their native host or landrace. ANOVA of the average diseased leaf area, which can be interpreted as the performance of a given *P. oryzae* isolate on a given accession, showed that the effect of the isolate*accession combination (F = 1.8, P < 10^-16^, df = 1088) remains highly significant after removing the significant effect of the experimental replicate (F = 2092.6, P < 10^-16^, df = 1). We used this ANOVA model to estimate the adjusted performance of each isolate on each accession. Heatmaps of the adjusted performance (Fig. 4) showed differential quantitative responses on the different rice accessions for all *P. oryzae* isolates, with only one isolate being very weakly aggressive (green color on Fig. 4) on all rice accessions (CH1877) and no isolate being highly aggressive (red color on Fig. 4) on all rice accessions. Except for four isolates (CH1897, CH1900, CH1901, CH1905), the adjusted performance of each *P. oryzae* isolate was not significantly better on its native rice accessions than on all other accessions (Fig. 4A, Fig. SI3.3). Hierarchical clustering by columns and lines (Fig. 4B) showed a lack of structure in the matrix, neither according to rice landraces, or to genetic lineages of *P. oryzae* isolates themselves. Finally, the average performance of all *P. oryzae* isolates originating from plants of a given landrace was not significantly better on all plants from this landrace than on plants of other landraces (Fig. 5). Altogether, these results strongly suggest that *P. oryzae* population in YYT did not adapt specifically to their native rice accessions or to any *indica* landrace.

**Figure 4.**
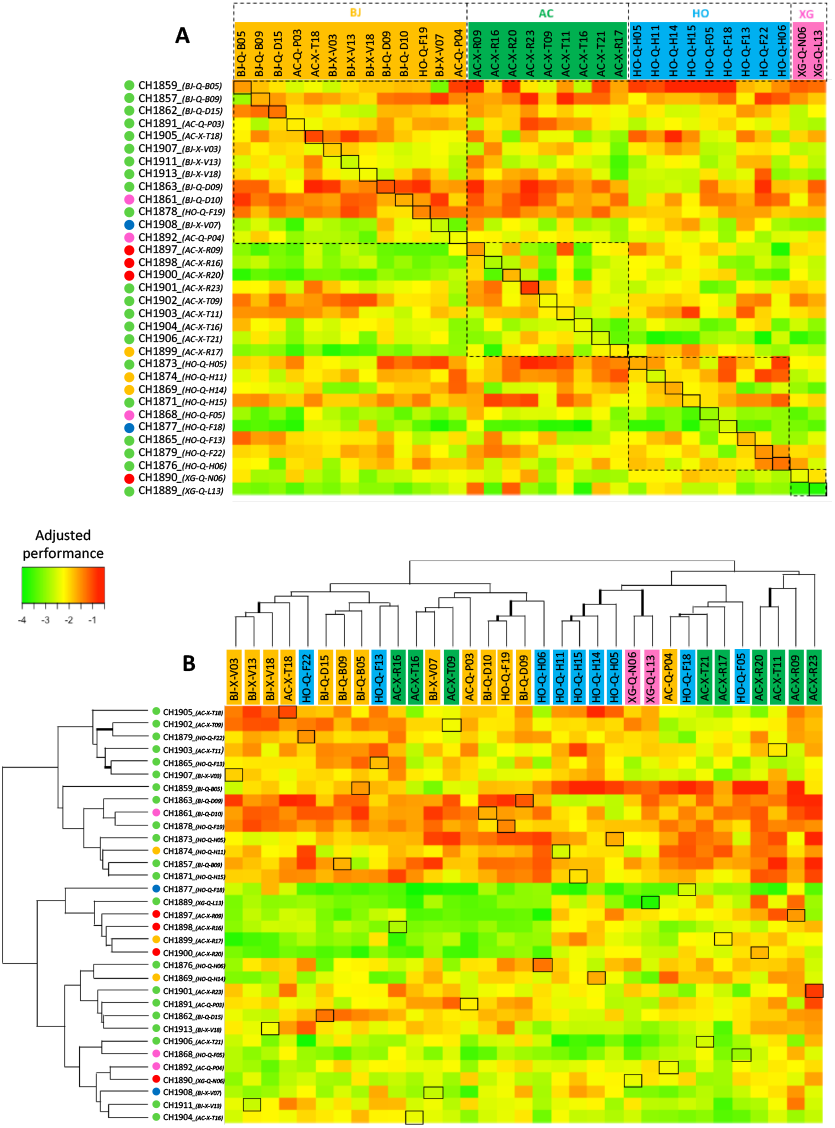
Heatmap of quantitative interactions between 33 *P. oryzae* isolates (by lines) and their corresponding rice accessions (by colomns). The plotted variable is the adjusted performance (log-transformation of the mean percentage of diseased leaf surface over 4 scored leaves, estimated from the ANOVA model) and varies from −4 (low performance) to 0 (high performance). Rice accessions are coloured according to the genetic cluster they belong to (according to clustering analysis of rice genotypes, see Fig. 3 left panel). Coloured dots besides *P. oryzae* isolates’ names correspond to the genetic lineage *P. oryzae* genotypes belong to (according to clustering analysis of *P. oryzae* genotypes, see Fig. 3 right panel); (green: lineage YYT1; red: lineage YYT2; pink: lineage YYT3; yellow: worldwide lineages 1 and 5; blue: worldwide lineage 3). A: Heatmap without hierarchical clustering; rice accessions are ordered by rice genetic clusters, *P. oryzae* isolates are ordered so that paired samples (framed) are along the diagonal. B: Heatmap with hierarchical clustering by lines and columns; branches with bootstrap support above 50 are bold (10,000 bootstrap replicates); paired samples are framed.

**Figure 5.**
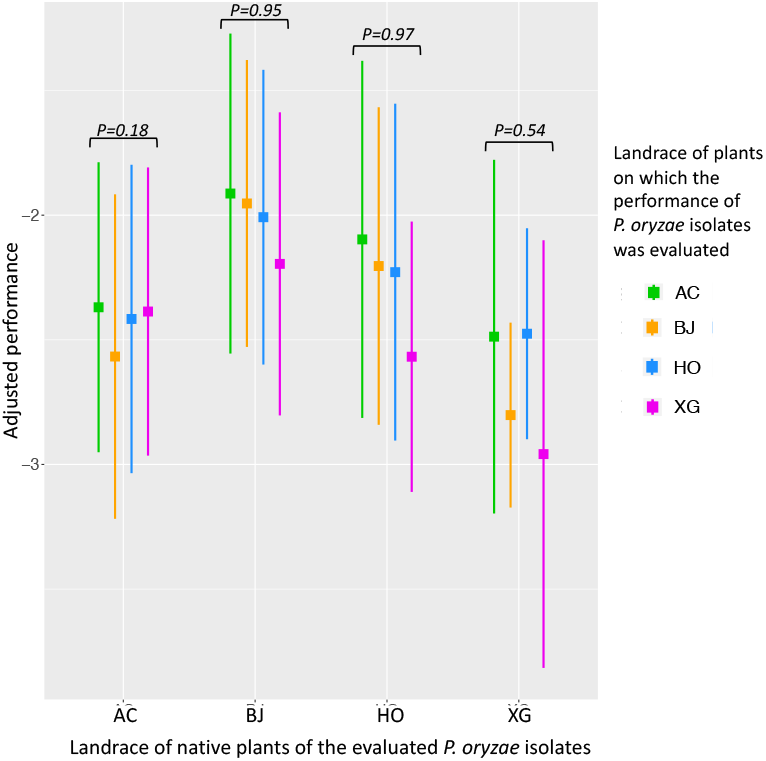
Boxplots of the average performance of *P. oryzae* isolates sampled on plants of from a given landrace, measured on plants of this landrace and of the other landraces. P-values of the corresponding contrasts are indicated above each set of boxplots.

## DISCUSSION

In this study we have characterized the genetic and pathotypic population structure of *P. oryzae* pathogens infecting traditional *indica* varieties in the Yuanyang terraces of rice paddies. This traditional agrosystem, maintained over centuries (He et al., 2011), and where rice disease pressure was reported to be low (Sheng, 1990), provides a unique opportunity to decipher the impact of crop diversity on disease epidemics, especially of rice blast.

Our first important observation based on analysis of microsatellite and whole genome genetic variation, is the finding of new lineages of the rice blast pathogen endemic to YYT. Our analysis of population structure based on microsatellite (513 isolates) and whole genome (46 isolates) datasets revealed multiple lineages of *P. oryzae* coexisting in YYT, with relatively high levels of standing variation compared to previous results from Gladieux *et al*. (2018b) and Saleh *et al*. (2014). Three genetic lineages endemic to YYT, coexisted with two of the four worldwide lineages previously described (Gladieux *et al*., 2018b). Although the global linkage disequilibrium inferred from microsatellite data was significantly different from 0, analyses of LD-decay, PHI-tests and reticulations within each of the five lineages provided contrasted information regarding the existence of recombination (Supplementary Information 2, Fig. SI2.2). Both mating types were found in sympatry (same village; data not shown) within the 46 YYT *P. oryzae* fully sequenced isolates, but they were found within the same lineage only for WL1 (Supplementary Information 2, Table SI2.3), which, together with significant PHI-test and reticulations observed for this lineage, was consistent with the fact that it has been described as recombinant (Gladieux et al., 2018; Latorre et al., 2020; Saleh et al., 2014; Thierry et al., 2021). Conversely, only Mat-2 isolates were found among the 30 fully sequenced isolates assigned to lineage YYT1. Therefore, significant PHI-test and reticulations observed in the minimum spanning network for this lineage could be due to scarce genetic exchanges among lineages (Gladieux et al., 2018), or to footprints of historical recombination. In invasive pathogens, higher genetic diversity and signatures of recombination are expected in older, source populations (Ali et al., 2014; Thach et al., 2016). Our observations therefore suggest that YYT area is very close to, if not included into, the centre of origin of rice-infecting *P. oryzae* pathogens in continental Southeast Asia hypothesized by previous studies (Gladieux et al., 2018; Saleh et al., 2012; Zhong et al., 2018).

Our second important observation is that *P. oryzae* populations are not specialized to traditional rice landraces in YYT. For pathogens mating within or onto their hosts, specialization should drastically restrict encounters of potential mates and reduce survival of offspring due to maladaptation of immigrants and hybrid offspring (Giraud et al., 2010; Gladieux et al., 2011), which should align the structure of pathogenic populations on that of the host. Our genotyping-by-sequencing data show that the rice accessions from YYT are structured into landraces with relatively high levels of genetic diversity, both within and among landraces, confirming previous findings based on 24 microsatellite markers (Gao et al., 2012). *P. oryzae* populations are also structured into different lineages, but our analysis reveals a complete lack of host-pathogen genetic co-structure. The fact that population subdivision in the pathogen does not mirror population subdivision in the host strongly suggests a lack of specialization to the host. We also analysed the phenotypic relationships among paired *P. oryzae* / rice samples by cross inoculating all isolates on their native and non-native plants. We showed that nearly all qualitative interactions were compatible, indicating that the all major resistance genes present in the YYT rice gene pools were defeated by the *P. oryzae* population. Quantitative interactions indicated that *P. oryzae* isolates did not perform significantly better on their native than on their non-native plants, and that *P. oryzae* isolates originating from plants of the same landrace did not perform significantly better on this landrace than on any other landraces, thus leading to rejection of the “home versus away” criteria of local adaptation (Blanquart et al., 2013). Together, the results of our analysis of population genetic and pathotypic structure therefore reveals a complete lack of *P. oryzae* specialization to rice landraces. Although our results are consistent with a generalist life style in *P. oryzae* population, maladaptation cannot be definitely ruled out. Maladaptation in pathogens describes the case where isolates performance is significantly better on non-native hosts than on native hosts (Kniskern et al., 2011). Given that we have assessed performance using an overall infection trait (the percentage of diseased leaf surface), we cannot exclude that other traits involved in fungal fitness that are not captured by our index (e.g. number of lesions, proportion of HR lesions among infective lesions, growth speed of lesion, sporulation capacity) would reveal such a maladaptation pattern.

Rice landraces and their *P. oryzae* pathogens from the Yuanyang terraces therefore represent a model system in which the pathogen is specialized to *indica* and *japonica* rice subspecies (Liao et al. 2016), but not specialized to the various landraces of *indica* rice. This is consistent with predictions that the nature of the mechanism underlying immunity in the host or avirulence in the pathogen and the magnitude of divergence in immune systems between hosts has an impact on the likelihood of pathogen specialization (Giraud et al., 2010; Schulze-Lefert and Panstruga, 2011). Liao *et al*. (2016) showed that in the YYT area where japonica and indica rice subspecies are cultivated in sympatry, differences in immune systems between *indica* and *japonica* subspecies would prevent the emergence of populations with a generalist lifestyle on both hosts, because a large effector complement is required to infect *japonica* rice, while *indica* rice has a larger repertoire of immune receptors and therefore greater capacity to detect effectors that will trigger immunity. Differences in repertoires of immune receptors among *indica* landraces might be sufficient to lead to the emergence of specialized *P. oryzae* populations in stable and homogeneous conditions. The maintenance of generalist *P. oryzae* genotypes highlighted by our results could be explained by the elevated heterogeneity of the “host landscape” in the YYT area, with spatio-temporal variation in rice genotypes distribution (elevated genetic diversity within and among rice landraces, spatial mosaics of paddy fields sown with different landraces, and temporal turnover of the mosaics) impeding the emergence of specialized pathogens on specific *indica* plant genotypes or landraces. Unlike populations of *P. oryzae* from modern agrosystems, which tend to be largely clonal and infecting relatively stable and homogenous host populations (Gladieux et al., 2018; Zhong et al., 2018), populations infecting YYT landraces displayed higher genotypic and genetic diversities with occurrence of recombination. This last feature may contribute to the maintenance of the generalist lifestyle by re-shuffling virulence alleles among *P. oryzae* pathogens. Finally, the Yuanyang Terraces bring to light an interesting paradox, in which the populations of the pathogen are diverse and recombinant - characteristics of populations with high adaptive potential - while they generate less yield losses than the clonal populations observed in other parts of the globe. This suggests that the epidemiological models that should help the re-engineering of agrosystems should not ignore the scenarios leading to the emergence of recombinant and diverse pathogenic populations.

## Supporting information

Summlemntal text S1

Summlemntal text S2

Summlemntal text S3

## Acknowledgements

We thank INRAe SMaCH Metaprogram for granting this study, and the International Associate Laboratory (LIA) Plantomix for financial and logistic support to field visits. The contribution of S.A. was supported by an AgreenSkills+ postdoctoral fellowship.

## Authors’ contributions

BJ, HH, HA, JBM, EF, XE and DT collected samples. SA, TD, PG, HA, IM, JM, SCA, BJ, DT, JBM and EF contributed to isolation of the pathogen and phenotyping experiments. SA, SCA, HA and JM conducted the molecular work. SA, PG, SR and EF conducted population genetics and genomic analyses of the data. TD contributed to microsatellite genotyping. FC and AL contributed to analyses of mating type in the genomic data. SA, FB, IM and EF conducted the image analyses and statistical analyses of phenotypic data. SA, PG, HH, JBM and EF wrote the manuscript. All authors contributed to the revision of the manuscript. SA, PG, HH, JBM and EF designed the study. DT, JBM and EF provided resources for the study.

