## Supplementary material for "Coevolution with Spatially Structured Rice Landraces Maintains Multiple Generalist Lineages in the Rice Blast Pathogen": Summlemntal text S1

**Ali et al. 2021. Supplementary Information SI2**

**SUPPORTING MATERIAL ON ANALYSES OF *P. ORYZAE* POPULATION STRUCTURE**

Surveillance was conducted in the rice cultivated area of Yuanyang terrace to collect samples of blast infected rice samples (Fig. SI2.1). A total of 513 isolates were collected and genotyped using microsatellite. 46 of these isolates were fully sequenced using Illumina short-reads technology.

**Fig. SI2.1. Localization of YYT area within the Yunnan province of China, and of the villages in which *P. oryzae* and rice were sampled.**

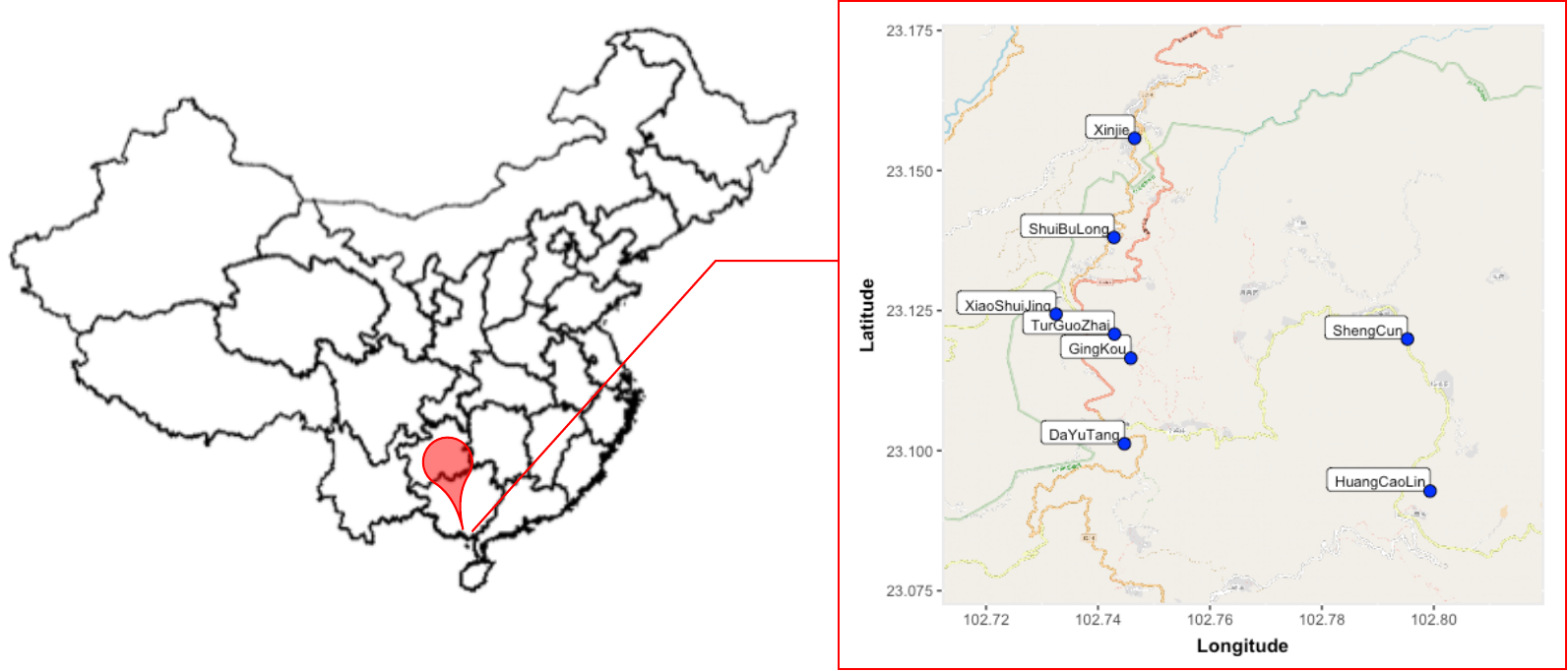

**Assignment of the 46 fully-sequenced isolates from YYT to different genetic lineages.**

We pooled whole-genome SNPs data from the 46 fully-sequenced isolates from YYT with whole-genome SNPs data previously acquired from 48 isolates representative of worldwide genetic diversity (Gladieux et al. 2018b). Assignment of YYT isolates to the different lineages was based on the phylogenetic network inferred using Splistree (Fig. 2C, Table SI2.1)

**Table SI2.1. Assignment of 46 *P. oryzae* isolates to genetic lineages using full genome SNPs data.** W-Lineages: worldwide lineages as previously described (Gladieux et al. 2018, Latorre et al. 2020, Thierry et al. 2021). Lineages YYT1 to YYT3: genetic lineage specific to YYT area.

| **Isolate** | **Variety** | **Lineage assigned** |
| --- | --- | --- |
| CH1869 | Hongjiao | W-Lineage 1 |
| CH1874 | Hongjiao | W-Lineage 1 |
| CH1899 | Acuce | W-Lineage 1 |
| CH1877 | Hongjiao | W-Lineage 3 |
| CH1882 | Hongyang | W-Lineage 3 |
| CH1885 | Hongyang | W-Lineage 3 |
| CH1893 | Acuce | W-Lineage 3 |
| CH1908 | Baijiao | W-Lineage 3 |
| CH1889 | Xiaogu | W-Lineage 1 |
| CH1857 | Baijiao | YYT1 |
| CH1858 | Baijiao | YYT1 |
| CH1862 | Baijiao | YYT1 |
| CH1863 | Baijiao | YYT1 |
| CH1865 | Hongjiao | YYT1 |
| CH1866 | Hongjiao | YYT1 |
| CH1871 | Hongjiao | YYT1 |
| CH1873 | Hongjiao | YYT1 |
| CH1876 | Hongjiao | YYT1 |
| CH1878 | Hongjiao | YYT1 |
| CH1879 | Hongjiao | YYT1 |
| CH1883 | Hongyang | YYT1 |
| CH1884 | Hongyang | YYT1 |
| CH1886 | Hongyang | YYT1 |
| CH1887 | Hongyang | YYT1 |
| CH1891 | Acuce | YYT1 |
| CH1895 | Acuce | YYT1 |
| CH1896 | Acuce | YYT1 |
| CH1901 | Acuce | YYT1 |
| CH1902 | Acuce | YYT1 |
| CH1903 | Acuce | YYT1 |
| CH1904 | Acuce | YYT1 |
| CH1905 | Acuce | YYT1 |
| CH1906 | Acuce | YYT1 |
| CH1907 | Baijiao | YYT1 |
| CH1910 | Baijiao | YYT1 |
| CH1911 | Baijiao | YYT1 |
| CH1913 | Baijiao | YYT1 |
| CH1915 | Baijiao | YYT1 |
| CH1859 | Baijiao | YYT1 |
| CH1890 | Xiaogu | YYT2 |
| CH1897 | Acuce | YYT2 |
| CH1898 | Acuce | YYT2 |
| CH1900 | Acuce | YYT2 |
| CH1861 | Baijiao | YYT3 |
| CH1868 | Hongjiao | YYT3 |
| CH1892 | Acuce | YYT3 |

**Analysis of recombination footprints in microsatellite and genomic *P. oryzae* population datasets.**

Signatures of recombination were searched using both microsatellite and genomic data. First, we used the microsatellite dataset to estimate the rate of sexual reproduction within the whole *P.oryzae* population using the CLONCASE method (Ali et al. 2016), which is based on resampling of multilocus microsatellite genotypes over years, assuming 10 clonal cycles before the onset of sexual cycle for each year. The estimated rates of sexual reproduction in the overall YYT *P. oryzae* population varied from 15 to 100 % according to the time period considered (Table SI2.2).

**Table SI2.2. Effective population size (Ne) and rate of sexual reproduction (s) of *P. oryzae*, estimated with the CLONCASE method based on re-sampling of multilocus microsatellite genotypes (MLGs) over years.**

| **Parameter** | **Time period** | | | | | |
| --- | --- | --- | --- | --- | --- | --- |
|  | **2009-11** | **2011-12** | **2012-13** | **2013-15** | **2015-16** | **2009-16** |
| **Ne (effective population size)** | 1859 | 473 | 91 | 8819 | 58 | 273 |
| **s (rate of sexual reproduction)** | 0.83 | 0.15 | 0.98 | 1.00 | 0.20 | 0.90 |
| **Number of different MLGs resampled** | 2 | 5 | 1 | 0 | 1 | 0 |

Signatures of recombination were also investigated within each of the five inferred genetic lineages using the genomic data retrieved from the complete resequencing of the 46 YYT *P. oryzae* isolates. LD decay along the genome was assessed within each genetic lineage using PopLDdecay (Zhang et al. 2019). We also used SPLITSREE to generate minimum spanning networks within each lineage, and to perform the Phi-test that assesses pairwise homoplasy and exchangeability of sites under the null hypothesis of no recombination (significant deviation from the null hypothesis is tested using random permutations of the positions of the SNPs). Reticulations were observed in the minimum spanning network for lineage YYT1, and to a lesser extent for lineages WL1, WL3 and YYT2. Although all PHI-tests were significant, none of the LD decay patterns showed significant evidence of recombination (Fig. SI2.3).

**Fig. SI2.2. Recombination among haplotypes identified using the full set of 66,102 SNPs without missing data from the whole-genome resequencing of 46 *P. oryzae* isolates. A. Linkage disequilibrium (LD, estimated using r2) against physical distance. B. Minimum spanning networks within each of the 5 *P. oryzae* lineages.** WL1 & WL3: Worldwide lineage-1 and Worldwide lineage-3. YYT1- YYT3 refers to the three genetic lineages specific to YYT.

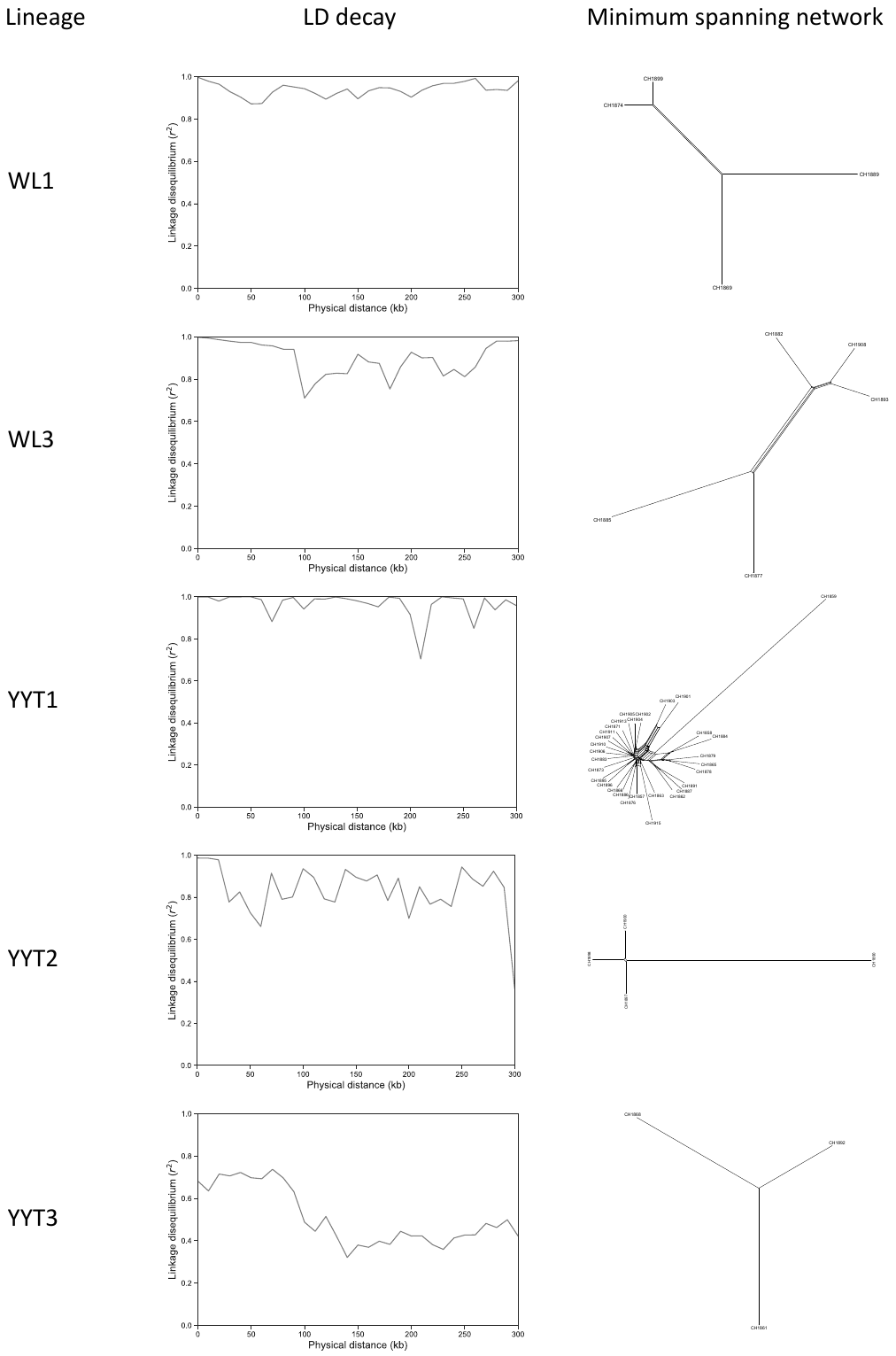

Finally, the presence of both mating types within the whole sample and within each lineage was assessed as a proxy for possibility of sexual reproduction (Saleh et al. 2012). For that, we searched for the Mat1.1 and Mat1.2 idiomorphs (corresponding to MAT1.2 and MAT1.1 in Kanamori 2007, respectively) in each of the 46 fully sequenced isolates using a BLAST procedure. For each isolate, raw Illumina data were assembled using ABySS 2.0 (Jackman 2017), by varying the k-mer from 20 to 90, and the assembly with the highest N50 value was selected. The selected assembly served as subject for BLASTn procedures with the sequences of Mat1.1 (3,736 bp) and Mat1.2 (4,650 bp) idiomorphs as queries, respectively. Although both mating types were present in the whole dataset in unbalanced proportions (87% Mat1.2 isolates, 13% Mat1.1 isolates), they were never found together within a single lineage expect for lineage WL1 (Table SI2.4).

**Table SI2.3.** **Assignment of 46 *P. oryzae* isolates to Mat1.1 or Mat1.2 mating types.** Sequences of Mat1.1 (3,736 bp) and Mat1.2 (4,650 bp) idiomorphs served as queries for BLASTn against the de-novo genomic assembly of each isolate. Genetic lineage to which each isolate belongs follows Splitstree (Fig. 2C) and DAPC clustering analysis (Fig. 3 right panel) of *P. oryzae* genotypes.

| **Isolate** | **Lineage** | **Identity Mat1.1 (%)** | **Coverage Mat1.1 (%)** | **Identity Mat1.2 (%)** | **Coverage Mat1.2 (%)** | **Mating type assignment** |
| --- | --- | --- | --- | --- | --- | --- |
| CH1869 | W-Lineage 1 | 34.13 | 34.39 | 99.91 | 99.93 | Mat1.2 |
| CH1874 | W-Lineage 1 | 99.87 | 100 | 27.44 | 27.63 | Mat1.1 |
| CH1899 | W-Lineage 1 | 99.87 | 100 | 27.44 | 27.63 | Mat1.1 |
| CH1889 | W-Lineage 1 | 99.87 | 100 | 27.44 | 27.63 | Mat1.1 |
| CH1877 | W-Lineage 3 | 34.13 | 34.39 | 99.57 | 99.59 | Mat1.2 |
| CH1882 | W-Lineage 3 | 34.13 | 34.39 | 99.49 | 99.51 | Mat1.2 |
| CH1885 | W-Lineage 3 | 34.13 | 34.39 | 99.49 | 99.51 | Mat1.2 |
| CH1893 | W-Lineage 3 | 34.13 | 34.39 | 99.91 | 99.93 | Mat1.2 |
| CH1908 | W-Lineage 3 | 34.13 | 34.39 | 99.91 | 99.93 | Mat1.2 |
| CH1890 | YYT2 | 34.13 | 34.39 | 99.91 | 99.93 | Mat1.2 |
| CH1897 | YYT2 | 34.13 | 34.39 | 99.57 | 99.59 | Mat1.2 |
| CH1898 | YYT2 | 34.13 | 34.39 | 99.91 | 99.93 | Mat1.2 |
| CH1900 | YYT2 | 34.13 | 34.39 | 99.57 | 99.59 | Mat1.2 |
| CH1861 | YYT3 | 99.87 | 100 | 27.44 | 27.63 | Mat1.1 |
| CH1868 | YYT3 | 99.87 | 100 | 27.44 | 27.63 | Mat1.1 |
| CH1892 | YYT3 | 99.87 | 100 | 27.44 | 27.63 | Mat1.1 |
| CH1857 | YYT1 | 34.13 | 34.39 | 99.89 | 99.93 | Mat1.2 |
| CH1858 | YYT1 | 34.13 | 34.39 | 99.55 | 99.59 | Mat1.2 |
| CH1859 | YYT1 | 34.13 | 34.39 | 99.91 | 99.93 | Mat1.2 |
| CH1863 | YYT1 | 34.13 | 34.39 | 99.89 | 99.93 | Mat1.2 |
| CH1865 | YYT1 | 34.13 | 34.39 | 99.89 | 99.93 | Mat1.2 |
| CH1866 | YYT1 | 34.13 | 34.39 | 99.89 | 99.93 | Mat1.2 |
| CH1873 | YYT1 | 34.13 | 34.39 | 99.55 | 99.59 | Mat1.2 |
| CH1878 | YYT1 | 34.13 | 34.39 | 99.89 | 99.93 | Mat1.2 |
| CH1879 | YYT1 | 34.13 | 34.39 | 99.89 | 99.93 | Mat1.2 |
| CH1883 | YYT1 | 34.13 | 34.39 | 99.55 | 99.59 | Mat1.2 |
| CH1884 | YYT1 | 34.13 | 34.39 | 99.55 | 99.59 | Mat1.2 |
| CH1886 | YYT1 | 34.13 | 34.39 | 99.89 | 99.93 | Mat1.2 |
| CH1887 | YYT1 | 34.13 | 34.39 | 99.89 | 99.93 | Mat1.2 |
| CH1895 | YYT1 | 34.13 | 34.39 | 99.55 | 99.59 | Mat1.2 |
| CH1896 | YYT1 | 34.13 | 34.39 | 99.89 | 99.93 | Mat1.2 |
| CH1901 | YYT1 | 34.13 | 34.39 | 99.55 | 99.59 | Mat1.2 |
| CH1902 | YYT1 | 34.13 | 34.39 | 99.89 | 99.93 | Mat1.2 |
| CH1904 | YYT1 | 34.13 | 34.39 | 99.89 | 99.93 | Mat1.2 |
| CH1906 | YYT1 | 34.13 | 34.39 | 99.89 | 99.93 | Mat1.2 |
| CH1910 | YYT1 | 34.13 | 34.39 | 99.89 | 99.93 | Mat1.2 |
| CH1871 | YYT1 | 34.13 | 34.39 | 99.89 | 99.93 | Mat1.2 |
| CH1876 | YYT1 | 34.13 | 34.39 | 99.89 | 99.93 | Mat1.2 |
| CH1891 | YYT1 | 34.13 | 34.39 | 99.89 | 99.93 | Mat1.2 |
| CH1915 | YYT1 | 34.13 | 34.39 | 99.89 | 99.93 | Mat1.2 |
| CH1862 | YYT1 | 34.13 | 34.39 | 99.89 | 99.93 | Mat1.2 |
| CH1905 | YYT1 | 34.13 | 34.39 | 99.55 | 99.59 | Mat1.2 |
| CH1907 | YYT1 | 34.13 | 34.39 | 99.89 | 99.93 | Mat1.2 |
| CH1903 | YYT1 | 34.13 | 34.39 | 99.89 | 99.93 | Mat1.2 |
| CH1913 | YYT1 | 34.13 | 34.39 | 99.89 | 99.93 | Mat1.2 |
| CH1911 | YYT1 | 34.13 | 34.39 | 99.89 | 99.93 | Mat1.2 |
