## Supplementary material for "Coevolution with Spatially Structured Rice Landraces Maintains Multiple Generalist Lineages in the Rice Blast Pathogen": Summlemntal text S2

**Ali et al. 2021. Supplementary Information SI3**

**SUPPORTING MATERIAL ON ANALYSES OF HOST-PATHOGEN INTERACTION**

Microsatellite genotyping of the isolates collected on various traditional landraces revealed a lack of any host specific structure in the studied population. Instead of prevalence of specific multilocus genotypes or lineages on a given host landrace, an overall high diversity was observed on various host landraces (Table SI3.1), with resampling of multilocus genotypes on several landraces (Table SI3.2).

**Table SI3.1. Genotypic diversity, gene diversity, and linkage disequilibrium inferred from microsatellite data of *P. oryzae* sampled on different host landraces in YYT.** Genotypic diversity was measured by Simpsons diversity and evenness indexes. All metrics were estimated with POPPR package. Only rice landraces for which *P. oryzae* sample size was over 5 are shown.

| **YYT rice ecotype** | **Rice landrace** | **Sample size** | **No of MLG** | **Simpson’s diversity** | **Evenness** | **Gene diversity** | **r_D_** |
| --- | --- | --- | --- | --- | --- | --- | --- |
| **Indica** | Acuce | 132 | 59 | 0.863 | 0.317 | 0.483 | 0.281 |
|  | Hongjiao | 36 | 36 | 0.972 | 1.000 | 0.626 | 0.068 |
|  | Cheran | 26 | 19 | 0.932 | 0.864 | 0.419 | 0.185 |
|  | Xiaogu | 25 | 22 | 0.947 | 0.909 | 0.562 | 0.713 |
|  | Zaogu | 24 | 18 | 0.934 | 0.907 | 0.490 | 0.913 |
|  | Chezuo | 22 | 12 | 0.868 | 0.762 | 0.393 | 0.456 |
|  | Baijiao | 21 | 20 | 0.948 | 0.974 | 0.376 | 0.223 |
|  | Lujiaogu | 14 | 14 | 0.929 | 1.000 | 0.605 | 0.309 |
|  | Heigu | 6 | 5 | 0.778 | 0.930 | 0.113 | 0.206 |
|  | Aizhegu | 7 | 7 | 0.857 | 1.000 | 0.503 | 0.534 |
|  | Chebu | 6 | 3 | 0.611 | 0.898 | 0.292 | 0.144 |
|  | Hongyang1 | 6 | 4 | 0.667 | 0.812 | 0.450 | 0.236 |
|  | Hongyang2 | 7 | 7 | 0.857 | 1.000 | 0.476 | 0.828 |
|  | Hongyang3 | 52 | 9 | 0.467 | 0.464 | 0.028 | 0.074 |
|  | Unknown | 15 | 2 | 0.391 | 0.817 | 0.000 | 1.000 |
| **Glutinous** | Huangpinuo | 23 | 18 | 0.922 | 0.812 | 0.273 | NA |
|  | Nuogu | 31 | 8 | 0.716 | 0.626 | 0.374 | 0.250 |
|  | Manchehonglue | 8 | 4 | 0.656 | 0.808 | 0.360 | 0.397 |

**Table SI3.2. Distribution of multilocus microsatellite genotypes (MLGs) of *P. oryzae* across multiple host landraces sampled in YYT.** The colour indicate the magnitude (from red: minimum value, to green: maximum value). All rice landraces were included here.

| ***P. oryzae* MLG** | **No. of hosts on which MLG was resampled** | **No. of isolates / MLG in overall population** |  | **Resampling according to host landrace** | | | | | | | | | | | | | | | | | | | | | | | | | |
| --- | --- | --- | --- | --- | --- | --- | --- | --- | --- | --- | --- | --- | --- | --- | --- | --- | --- | --- | --- | --- | --- | --- | --- | --- | --- | --- | --- | --- | --- |
|  |  |  |  | **YYT indica landraces** | | | | | | | | | | | | | | | | | | | | | **YYT glutinous landraces** | | | | |
|  |  |  |  | **Acuce** | **Aizhegu** | **Azheche** | **BaiJiao** | **Chebu** | **Cheran** | **Chezuo** | **unknown 1** | **Heigu** | **Hongjiao** | **Lengshuigu** | **Lujiaogu** | **Manchehonglue** | **unknown 2** | **Meiziche** | **Xiaogu** | **Zaoggu** | **Hongyang1** | **Hongyang2** | **Hongyang3** | **unknown 3** | **NuoGu** | **ZiNuo** | **HongNuo** | **HuaGu** | **HuangPiNuo** |
| MLG-210 | 10 | 109 |  | 43 |  |  | 1 | 3 | 2 | 6 |  |  |  |  |  | 2 |  |  |  | 3 |  |  | 37 | 11 |  | 1 |  |  |  |
| MLG-272 | 4 | 9 |  | 3 |  |  |  |  | 3 | 2 |  |  |  |  |  | 1 |  |  |  |  |  |  |  |  |  |  |  |  |  |
| MLG-205 | 4 | 5 |  | 1 | 1 | 1 |  |  |  |  |  |  |  |  |  |  |  |  |  | 2 |  |  |  |  |  |  |  |  |  |
| MLG-211 | 3 | 6 |  | 1 |  |  |  |  |  |  |  |  |  |  |  |  |  |  |  |  |  |  | 1 | 4 |  |  |  |  |  |
| MLG-203 | 3 | 4 |  | 2 |  |  |  |  |  |  |  |  |  |  | 1 |  |  |  |  | 1 |  |  |  |  |  |  |  |  |  |
| MLG-10 | 2 | 15 |  |  |  |  |  |  |  |  | 14 |  |  |  |  |  |  |  |  |  |  |  |  |  | 1 |  |  |  |  |
| MLG-162 | 2 | 13 |  | 5 |  |  |  |  |  |  |  |  |  |  |  |  |  |  |  |  |  |  | 8 |  |  |  |  |  |  |
| MLG-269 | 2 | 9 |  |  |  |  |  |  | 5 |  |  |  |  |  |  | 4 |  |  |  |  |  |  |  |  |  |  |  |  |  |
| MLG-58 | 1 | 8 |  |  |  |  |  |  |  |  |  |  |  |  |  |  |  |  |  |  |  |  |  |  | 4 |  |  |  | 4 |
| MLG-201 | 2 | 5 |  |  |  |  | 2 |  |  |  |  |  |  |  |  |  |  |  | 3 |  |  |  |  |  |  |  |  |  |  |
| MLG-66 | 2 | 5 |  | 2 |  |  |  |  |  |  |  |  |  |  |  |  |  | 3 |  |  |  |  |  |  |  |  |  |  |  |
| MLG-219 | 2 | 4 |  | 1 |  |  |  |  |  | 3 |  |  |  |  |  |  |  |  |  |  |  |  |  |  |  |  |  |  |  |
| MLG-265 | 2 | 4 |  |  |  |  |  |  | 3 |  |  |  |  |  |  |  |  |  |  | 1 |  |  |  |  |  |  |  |  |  |
| MLG-62 | 1 | 4 |  |  | 1 |  |  |  |  |  |  |  |  |  |  |  |  |  |  |  |  |  |  |  |  |  |  |  | 3 |
| MLG-200 | 2 | 3 |  |  |  |  |  |  |  |  |  |  | 1 |  |  |  |  |  | 2 |  |  |  |  |  |  |  |  |  |  |
| MLG-252 | 2 | 3 |  |  |  |  |  |  | 1 | 2 |  |  |  |  |  |  |  |  |  |  |  |  |  |  |  |  |  |  |  |
| MLG-134 | 2 | 2 |  | 1 |  |  |  |  |  |  |  |  |  |  |  |  |  |  |  |  |  |  |  |  |  | 1 |  |  |  |
| MLG-189 | 2 | 2 |  |  |  |  |  |  |  |  |  |  |  |  | 1 |  |  |  |  |  |  |  | 1 |  |  |  |  |  |  |
| MLG-206 | 2 | 2 |  | 1 |  |  |  |  | 1 |  |  |  |  |  |  |  |  |  |  |  |  |  |  |  |  |  |  |  |  |
| MLG-213 | 1 | 2 |  |  |  |  |  |  |  |  |  |  |  |  |  |  |  |  |  | 1 |  |  |  |  |  |  |  | 1 |  |
| MLG-214 | 2 | 2 |  |  |  |  |  |  |  | 1 |  |  |  |  |  |  |  |  |  |  | 1 |  |  |  |  |  |  |  |  |
| MLG-22 | 2 | 2 |  |  |  |  |  |  |  |  | 1 |  |  |  |  |  | 1 |  |  |  |  |  |  |  |  |  |  |  |  |
| MLG-221 | 2 | 2 |  |  |  |  |  |  |  |  |  |  |  |  | 1 |  |  |  | 1 |  |  |  |  |  |  |  |  |  |  |
| MLG-223 | 2 | 2 |  |  |  |  |  |  |  |  |  |  |  |  | 1 |  |  |  | 1 |  |  |  |  |  |  |  |  |  |  |
| MLG-242 | 1 | 2 |  |  |  |  |  |  |  |  |  |  |  |  |  |  |  |  |  | 1 |  |  |  |  |  |  |  | 1 |  |
| MLG-243 | 2 | 2 |  |  |  | 1 |  |  |  |  |  |  |  |  | 1 |  |  |  |  |  |  |  |  |  |  |  |  |  |  |
| MLG-244 | 2 | 2 |  | 1 |  |  |  |  |  |  |  |  |  |  |  |  |  |  | 1 |  |  |  |  |  |  |  |  |  |  |
| MLG-275 | 2 | 2 |  |  |  |  |  |  | 1 |  |  |  |  |  |  |  |  |  |  | 1 |  |  |  |  |  |  |  |  |  |
| MLG-284 | 2 | 2 |  |  |  |  |  |  | 1 |  |  |  |  |  |  |  |  |  |  |  |  |  |  |  |  | 1 |  |  |  |
| MLG-65 | 1 | 2 |  |  | 1 |  |  |  |  |  |  |  |  |  |  |  |  |  |  |  |  |  |  |  |  |  |  |  | 1 |
| MLG-94 | 1 | 2 |  |  | 1 |  |  |  |  |  |  |  |  |  |  |  |  |  |  |  |  |  |  |  |  |  |  |  | 1 |
| MLG-248 | 1 | 21 |  | 21 |  |  |  |  |  |  |  |  |  |  |  |  |  |  |  |  |  |  |  |  |  |  |  |  |  |
| MLG-51 | 1 | 13 |  |  |  |  |  |  |  |  |  |  |  |  |  |  | 13 |  |  |  |  |  |  |  |  |  |  |  |  |
| MLG-75 | 1 | 12 |  |  |  |  |  |  |  |  |  |  |  |  |  |  | 12 |  |  |  |  |  |  |  |  |  |  |  |  |
| MLG-39 | 1 | 4 |  |  |  |  |  |  |  |  | 4 |  |  |  |  |  |  |  |  |  |  |  |  |  |  |  |  |  |  |
| MLG-100 | 1 | 3 |  |  |  |  |  |  |  |  |  |  |  |  |  |  |  |  |  |  | 3 |  |  |  |  |  |  |  |  |
| MLG-45 | 1 | 3 |  |  |  |  |  |  |  |  |  |  |  |  |  |  |  |  |  |  |  |  |  |  | 3 |  |  |  |  |
| MLG-55 | 0 | 3 |  |  |  |  |  |  |  |  |  |  |  |  |  |  |  |  |  |  |  |  |  |  |  |  | 3 |  |  |
| MLG-11 | 1 | 2 |  |  |  |  |  |  |  |  | 2 |  |  |  |  |  |  |  |  |  |  |  |  |  |  |  |  |  |  |
| MLG-142 | 1 | 2 |  |  |  |  |  |  |  | 2 |  |  |  |  |  |  |  |  |  |  |  |  |  |  |  |  |  |  |  |
| MLG-154 | 1 | 2 |  | 2 |  |  |  |  |  |  |  |  |  |  |  |  |  |  |  |  |  |  |  |  |  |  |  |  |  |
| MLG-164 | 1 | 2 |  |  |  |  |  |  |  |  |  |  |  | 2 |  |  |  |  |  |  |  |  |  |  |  |  |  |  |  |
| MLG-167 | 1 | 2 |  |  |  |  |  |  |  |  |  |  |  |  |  |  |  |  |  | 2 |  |  |  |  |  |  |  |  |  |
| MLG-175 | 1 | 2 |  |  |  |  |  |  |  |  |  | 2 |  |  |  |  |  |  |  |  |  |  |  |  |  |  |  |  |  |
| MLG-199 | 1 | 2 |  | 2 |  |  |  |  |  |  |  |  |  |  |  |  |  |  |  |  |  |  |  |  |  |  |  |  |  |
| MLG-236 | 1 | 2 |  |  |  |  |  | 2 |  |  |  |  |  |  |  |  |  |  |  |  |  |  |  |  |  |  |  |  |  |
| MLG-253 | 1 | 2 |  |  |  |  |  |  |  |  |  |  |  |  |  |  |  |  |  | 2 |  |  |  |  |  |  |  |  |  |
| MLG-263 | 1 | 2 |  |  |  |  |  |  |  |  |  |  |  |  |  |  |  |  |  | 2 |  |  |  |  |  |  |  |  |  |
| MLG-99 | 1 | 2 |  | 2 |  |  |  |  |  |  |  |  |  |  |  |  |  |  |  |  |  |  |  |  |  |  |  |  |  |
| **Sample size** |  | **557** |  | **132** | **7** | **2** | **21** | **6** | **29** | **22** | **22** | **6** | **36** | **4** | **14** | **8** | **28** | **3** | **25** | **24** | **6** | **7** | **52** | **15** | **9** | **4** | **4** | **3** | **23** |

Phenotyping was done to assess the qualitative interactions between 45 *P. oryzae* isolates and their corresponding rice accessions (Fig. SI3.1). Maratelli and Co39 were used as positive control of compatible interactions. For each rice**P. oryzae* combination, compatible and incompatible interactions were scored on 6 to 8 leaves; the plotted values correspond to the percentage of compatible or incompatible interactions on the scored leaves. The overall experiment was repeated two independent times and revealed a high repeatability (Fig. SI3.2).

**Fig. SI3.1. Qualitative interactions between 45 *P. oryzae* isolates (by lines) and their corresponding rice accessions (by columns).** Rice accessions are colored according to the genetic cluster they belong to (according to clustering analysis of rice genotypes, see Fig. SI3.1 left panel), and *P. oryzae* isolates are colored according to the genetic group their native plants belong to (orange: BJ, green: AC, blue: HO, pink: XG). Rice accessions are ordered by rice genetic clusters, *P. oryzae* isolates are ordered so that paired samples (framed) are along the diagonal.

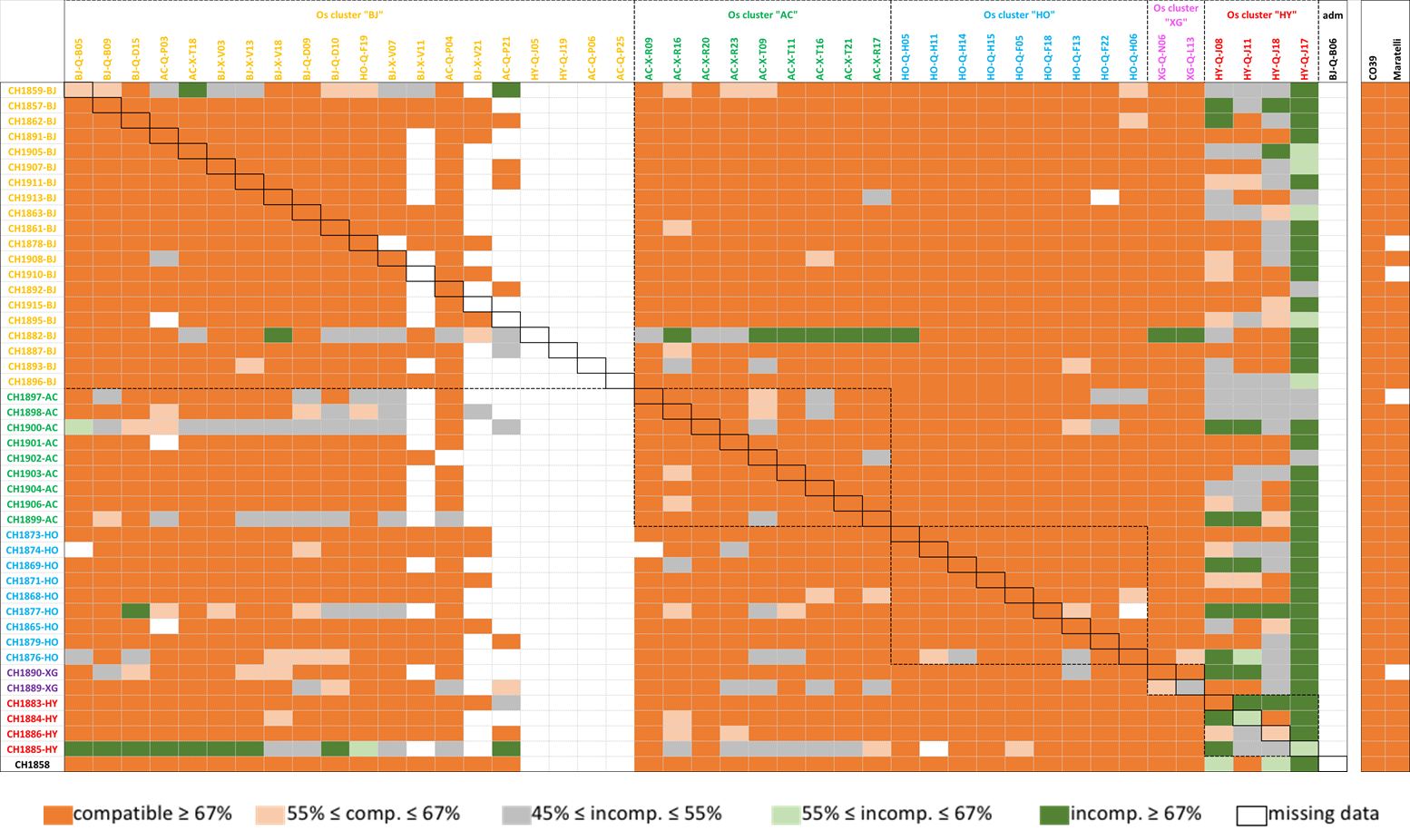

**Fig. SI3.2. Correlation between scoring of two independent replications of the cross-inoculation experiment of *P. oryzae* on rice host lines.**

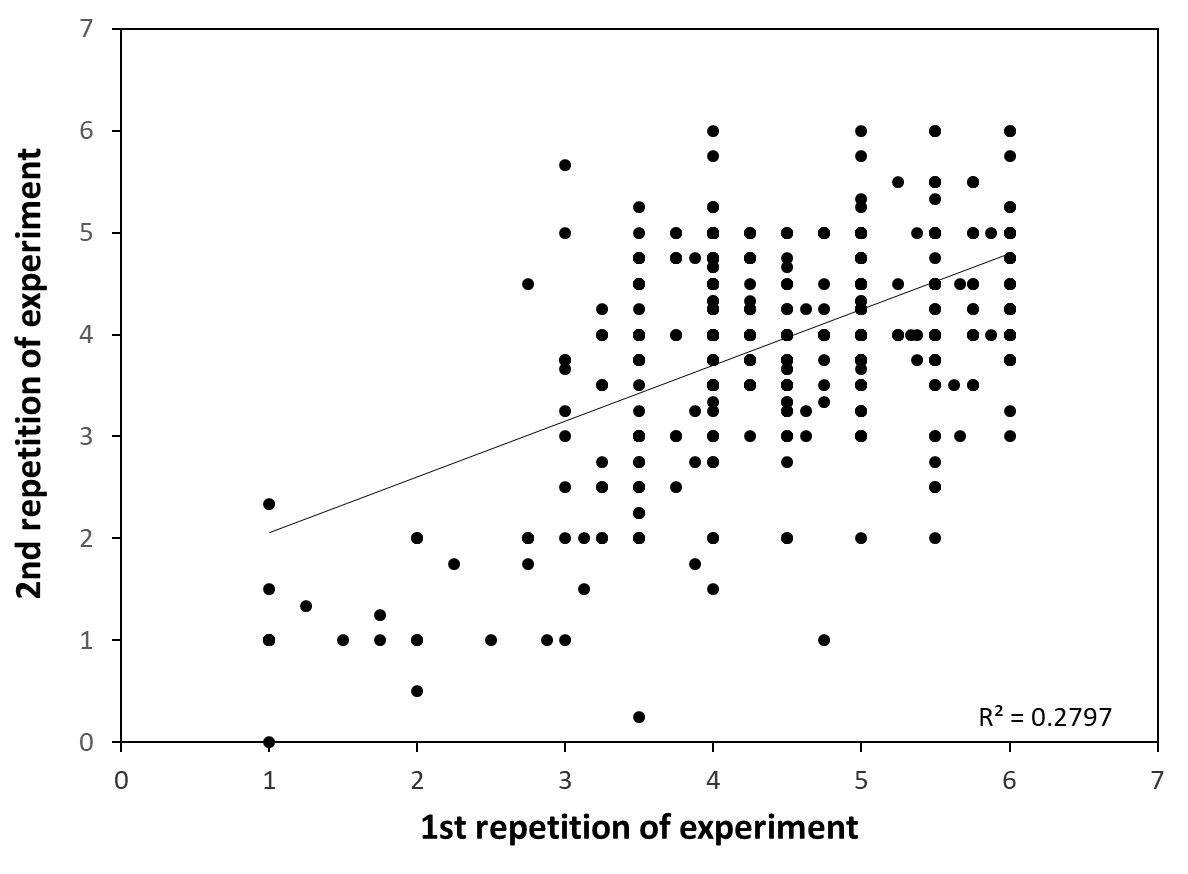

**Fig. S3.3. Adjusted performance of each *P. oryzae* isolate on its native plant (red dot) and on all other non-native plants (mean value = square, +- standard deviation).** Stars indicate significant contrasts (1 star: P<0.05, 2 stars: P<0.01). *P. oryzae* isolates are colored according to the genetic group their native plants belong to (orange: BJ, green: AC, blue: HO, pink: XG).

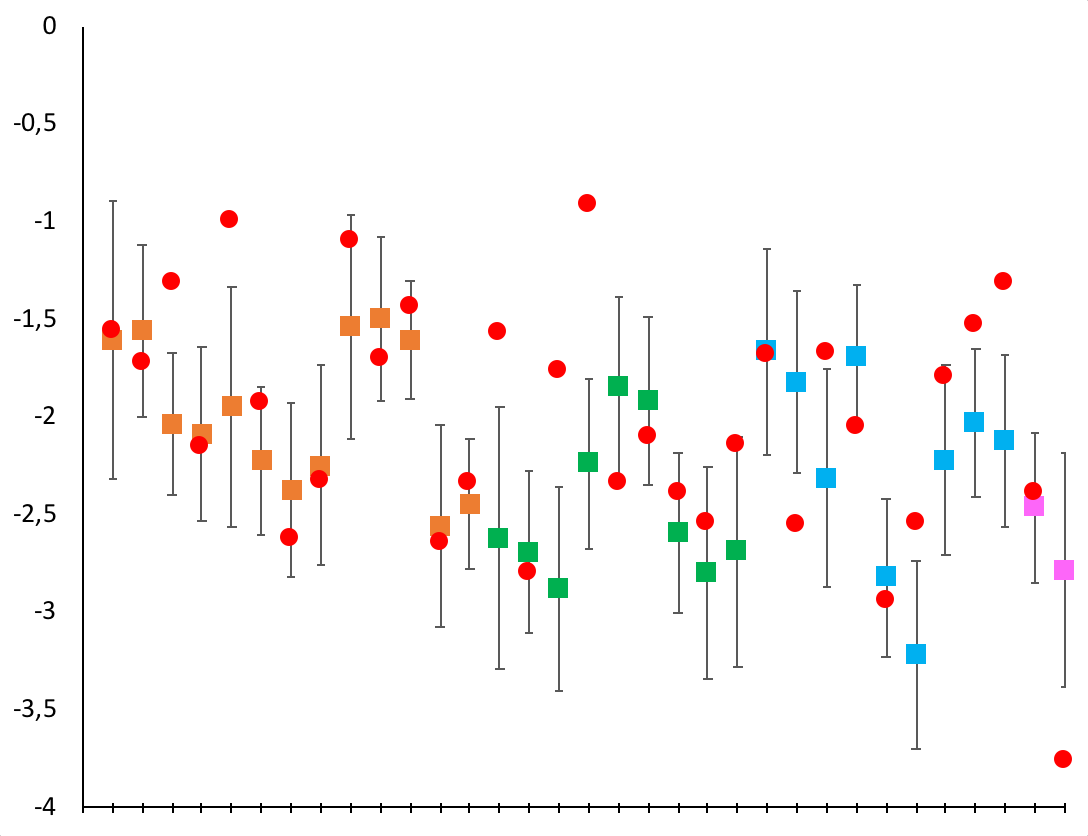

CH1859-BJ

CH1857-BJ

CH1862-BJ

CH1891-BJ

CH1905-BJ

CH1907-BJ

CH1911-BJ

CH1913-BJ

CH1863-BJ

CH1861-BJ

CH1878-BJ

CH1908-BJ

CH1892-BJ

CH1897-AC

CH1898-AC

CH1900-AC

CH1901-AC

CH1902-AC

CH1903-AC

CH1904-AC

CH1906-AC

CH1899-AC

CH1873-HO

CH1874-HO

CH1869-HO

CH1871-HO

CH1868-HO

CH1877-HO

CH1865-HO

CH1879-HO

CH1876-HO

CH1890-XJ

CH1889-XJ

*

*

*

*

*

*

*

*
