## Supplementary material for "Coevolution with Spatially Structured Rice Landraces Maintains Multiple Generalist Lineages in the Rice Blast Pathogen": Summlemntal text S3

**Ali et al. 2021. Supplementary Information SI1**

**SUPPORTING MATERIAL ON ANALYSES OF RICE LANDRACES**

**Phylogenetic analysis of the 92 rice accessions sampled in YYT together with 216 worldwide rice accessions.**

To assess the phylogenetic relationship of rice landraces from the Yuanyang terrace, China, a set of 92 rice accessions were selected for genotyping by sequencing (Table SI1.1). This set included the major cultivated landraces of the region, among which representatives of modern improved varieties, and representing different locations.

**Table SI1.1. List of the 92 YYT rice accessions analysed for population subdivision, with the vernacular name of the variety and the village in which they were sampled.**

| **Rice landrace code** | **GBS/Vcf Code** | **Year** | **landrace vernacular name** | **Location** | **Corresponding P. oryzae isolate** |
| --- | --- | --- | --- | --- | --- |
| YYT_Bai_B05 | B05 | 2015 | Baijiao | Jingkou | CH1859 |
| YYT_AC_B06 | B06 | 2015 | Baijiao | Jingkou | CH1858 |
| YYT_Bai_B09 | B09 | 2015 | Baijiao | Jingkou | CH1857 |
| YYT_Bai_D09 | D09 | 2015 | Baijiao | Jingkou | CH1863 |
| YYT_Bai_D10 | D10 | 2015 | Baijiao | Jingkou | CH1861 |
| YYT_Bai_D15 | D15 | 2015 | Baijiao | Jingkou | CH1862 |
| YYT_HO_E1 | E1 | 2015 | Hongjiao | Jingkou | - |
| YYT_HO_E11 | E11 | 2015 | Hongjiao | Jingkou | - |
| YYT_HO_E3 | E3 | 2015 | Hongjiao | Jingkou | - |
| YYT_HO_E5 | E5 | 2015 | Hongjiao | Jingkou | - |
| YYT_HO_E7 | E7 | 2015 | Hongjiao | Jingkou | - |
| YYT_HO_E9 | E9 | 2015 | Hongjiao | Jingkou | - |
| YYT_HO_F05 | F05 | 2015 | Hongjiao | Jingkou | CH1868 |
| YYT_HO_F13 | F13 | 2015 | Hongjiao | Jingkou | CH1865 |
| YYT_HO_F16 | F16 | 2015 | Hongjiao | Jingkou | CH1866 |
| YYT_HO_F18 | F18 | 2015 | Hongjiao | Jingkou | CH1877 |
| YYT_Bai_F19 | F19 | 2015 | Hongjiao | Jingkou | CH1878 |
| YYT_HO_F22 | F22 | 2015 | Hongjiao | Jingkou | CH1879 |
| YYT_HO_G1 | G1 | 2015 | Hongjiao | Jingkou | - |
| YYT_HO_G13 | G13 | 2015 | Hongjiao | Jingkou | - |
| YYT_HO_G3 | G3 | 2015 | Hongjiao | Jingkou | - |
| YYT_HO_G5 | G5 | 2015 | Hongjiao | Jingkou | - |
| YYT_HO_G7 | G7 | 2015 | Hongjiao | Jingkou | - |
| YYT_HO_G9 | G9 | 2015 | Hongjiao | Jingkou | - |
| YYT_HO_H05 | H05 | 2015 | Hongjiao | Jingkou | CH1873 |
| YYT_HO_H06 | H06 | 2015 | Hongjiao | Jingkou | CH1876 |
| YYT_HO_H11 | H11 | 2015 | Hongjiao | Jingkou | CH1874 |
| YYT_HO_H14 | H14 | 2015 | Hongjiao | Jingkou | CH1869 |
| YYT_HO_H15 | H15 | 2015 | Hongjiao | Jingkou | CH1871 |
| YYT_HO2_J05 | J05 | 2015 | Hongyang | Jingkou | CH1882 |
| YYT_HO2_J08 | J08 | 2015 | Hongyang | Jingkou | CH1883 |
| YYT_HO2_J11 | J11 | 2015 | Hongyang | Jingkou | CH1884 |
| YYT_HO2_J17 | J17 | 2015 | Hongyang | Jingkou | CH1885 |
| YYT_HO2_J18 | J18 | 2015 | Hongyang | Jingkou | CH1886 |
| YYT_HO2_J19 | J19 | 2015 | Hongyang | Jingkou | CH1887 |
| YYT_Xi_K1 | K1 | 2015 | Xiaogu | Jingkou | - |
| YYT_Xi_K4 | K4 | 2015 | Xiaogu | Jingkou | - |
| YYT_Xi_K8 | K8 | 2015 | Xiaogu | Jingkou | - |
| YYT_Xi_L13 | L13 | 2015 | Xiaogu | Jingkou | CH1889 |
| YYT_Xi_M1 | M1 | 2015 | Xiaogu | Jingkou | - |
| YYT_Xi_M10 | M10 | 2015 | Xiaogu | Jingkou | - |
| YYT_Xi_M4 | M4 | 2015 | Xiaogu | Jingkou | - |
| YYT_Xi_N06 | N06 | 2015 | Xiaogu | Jingkou | CH1890 |
| YYT_Bai_P03 | P03 | 2015 | Acuce | Jingkou | CH1891 |
| YYT_Bai_P04 | P04 | 2015 | Acuce | Jingkou | CH1892 |
| YYT_Bai_P06 | P06 | 2015 | Acuce | Jingkou | CH1893 |
| YYT_Bai_P21 | P21 | 2015 | Acuce | Jingkou | CH1895 |
| YYT_Bai_P25 | P25 | 2015 | Acuce | Jingkou | CH1896 |
| YYT_AC_Q1 | Q1 | 2015 | Acuce | Xiaoshuijing | - |
| YYT_AC_Q10 | Q10 | 2015 | Acuce | Xiaoshuijing | - |
| YYT_AC_Q12 | Q12 | 2015 | Acuce | Xiaoshuijing | - |
| YYT_AC_Q15 | Q15 | 2015 | Acuce | Xiaoshuijing | - |
| YYT_AC_Q16 | Q16 | 2015 | Acuce | Xiaoshuijing | - |
| YYT_AC_Q4 | Q4 | 2015 | Acuce | Xiaoshuijing | - |
| YYT_AC_Q7 | Q7 | 2015 | Acuce | Xiaoshuijing | - |
| YYT_AC_R09 | R09 | 2015 | Acuce | Xiaoshuijing | CH1897 |
| YYT_AC_R16 | R16 | 2015 | Acuce | Xiaoshuijing | CH1898 |
| YYT_AC_R17 | R17 | 2015 | Acuce | Xiaoshuijing | CH1899 |
| YYT_AC_R20 | R20 | 2015 | Acuce | Xiaoshuijing | CH1900 |
| YYT_AC_R23 | R23 | 2015 | Acuce | Xiaoshuijing | CH1901 |
| YYT_AC_S1 | S1 | 2015 | Acuce | Xiaoshuijing | - |
| YYT_AC_S10 | S10 | 2015 | Acuce | Xiaoshuijing | - |
| YYT_AC_S13 | S13 | 2015 | Acuce | Xiaoshuijing | - |
| YYT_AC_S15 | S15 | 2015 | Acuce | Xiaoshuijing | - |
| YYT_AC_S3 | S3 | 2015 | Acuce | Xiaoshuijing | - |
| YYT_AC_S6 | S6 | 2015 | Acuce | Xiaoshuijing | - |
| YYT_AC_S8 | S8 | 2015 | Acuce | Xiaoshuijing | - |
| YYT_AC_T09 | T09 | 2015 | Acuce | Xiaoshuijing | CH1902 |
| YYT_AC_T11 | T11 | 2015 | Acuce | Xiaoshuijing | CH1903 |
| YYT_AC_T16 | T16 | 2015 | Acuce | Xiaoshuijing | CH1904 |
| YYT_AC_T18 | T18 | 2015 | Acuce | Xiaoshuijing | CH1905 |
| YYT_AC_T21 | T21 | 2015 | Acuce | Xiaoshuijing | CH1906 |
| YYT_Bai_V03 | V03 | 2015 | Baijiao | Xiaoshuijing | CH1907 |
| YYT_Bai_V07 | V07 | 2015 | Baijiao | Xiaoshuijing | CH1908 |
| YYT_Bai_V11 | V11 | 2015 | Baijiao | Xiaoshuijing | CH1910 |
| YYT_Bai_V13 | V13 | 2015 | Baijiao | Xiaoshuijing | CH1911 |
| YYT_Bai_V18 | V18 | 2015 | Baijiao | Xiaoshuijing | CH1913 |
| YYT_Bai_V21 | V21 | 2015 | Baijiao | Xiaoshuijing | CH1915 |
| YYT_Hu_YYT27B | YYT27B | 2014 | Huangpinuo | Jingkou | - |
| YYT_Hu_YYT27R | YYT27R | 2014 | Huangpinuo | Jingkou | - |
| YYT_Bag_YYT31 | YYT31 | 2014 | Baijiao | Jingkou | - |
| YYT_Xi_YYT55 | YYT55 | 2014 | Xiaogu | Jingkou | - |
| YYT_Xi_YYT56R | YYT56R | 2014 | Xiaogu | Jingkou | - |
| YYT_Xi_YYT60 | YYT60 | 2014 | Xiaogu | Jingkou | - |
| YYT_Zn_YYT72 | YYT72 | 2014 | Zinuo | Jingkou | - |
| YYT_AC_YYT73 | YYT73 | 2014 | Acuce | Jingkou | - |
| YYT_Hu_YYT76B | YYT76B | 2014 | Huangpinuo | Jingkou | - |
| YYT_Hu_YYT76N | YYT76N | 2014 | Huangpinuo | Jingkou | - |
| YYT_Hu_YYT76R | YYT76R | 2014 | Huangpinuo | Jingkou | - |
| YYT_Hu_YYT77B | YYT77B | 2014 | Huangpinuo | Jingkou | - |
| YYT_Nu_YYT78 | YYT78 | 2014 | Nuogu | Jingkou | - |
| YYT_Bag_YYT80 | YYT80 | 2014 | Baijiao | Jingkou | - |

To compare YYT rice diversity to worldwide rice diversity, a first set of 1415 reference accessions publicly available (Huang et al., 2012; Wang et al. 2017) were pooled with the 92 YYT rice accessions to perform mapping against the rice reference genome IRGSP-1.0 Nipponbare (Kawahara et al. 2013) and SNP calling. This file contained a total of 1511 rice accessions and 7,827,056 positions. To remove positions and accessions with too many missing data, we proceeded in two steps. First, we checked the relative frequency of missing data in the worldwide reference accessions only, using in-house scripts, by varying the cutoff of relative frequency of missing data from 40% to 60%. The proportions of worldwide rice accessions having missing data were 98%, 92% and 83% for cutoffs of 40%, 50% and 60%, respectively (Table SI1.2). For further analyses, we kept only worldwide accessions with less than 60% missing SNP sites, and also removed accessions not assigned to any group by previous studies (admixed or intermediate). The resulting dataset contained a total of 308 rice accessions (92 YYT accessions + 216 worldwide accessions, Table SI1.3) and was then filtered to keep positions having less than 30% missing data, giving a total of 44,855 SNPs.

**Table** **SI1.2. Number of worldwide accessions to be retained after keeping accessions with different cut-off values for missing data (40%-60%).** * Accessions not assigned to any group in previous studies (Admixed and Intermedia) were also excluded.

| **Ecotype** | **Initially set of accessions** | **Cutoff-40%** | **Cutoff-50%** | **Cutoff-60%** | **Finally retained (Cutoff-60%)*** |
| --- | --- | --- | --- | --- | --- |
| Aromatic | 9 | 0 | 4 | 5 | 5 |
| AUS | 40 | 0 | 11 | 23 | 23 |
| Indica | 465 | 4 | 20 | 48 | 48 |
| Temperate japonica | 319 | 1 | 14 | 25 | 25 |
| Tropical japonica | 57 | 1 | 10 | 25 | 25 |
| *O. barthii* | 1 | 1 | 1 | 1 | 1 |
| *O. glaberrima* | 1 | 1 | 1 | 1 | 1 |
| *O. longiglumis* | 2 | 0 | 0 | 0 | 0 |
| *O. meridionalis* | 12 | 0 | 0 | 0 | 0 |
| *O. rufipogon-I* | 155 | 7 | 18 | 44 | 44 |
| *O. rufipogon-II* | 121 | 5 | 13 | 28 | 28 |
| *O. rufipogon-III* | 169 | 5 | 6 | 16 | 16 |
| Admixture | 36 | 2 | 10 | 23 | 0 |
| Intermedia | 28 | 1 | 1 | 1 | 0 |
| **Total** | **1415** | **28** | **109** | **240** | **216** |

**Table SI1.3. Number of rice accessions selected to assess the relationship of YYT accessions within representatives of the worldwide rice diversity.**

| **Ecotype** | **Huang et al. (2012)** | **Wang et al., (2017)** | **This study** | **Total** |
| --- | --- | --- | --- | --- |
| **Aromatic** | 3 | 2 | - | 5 |
| **AUS** | 1 | 22 | - | 23 |
| **Indica** | 13 | 35 | - | 48 |
| **Temperate Japonica** | 2 | 23 | - | 25 |
| **Tropical Japonica** | 4 | 21 | - | 25 |
| ***O. barti*** | 1 | - | - | 1 |
| ***O. glab*** | 1 | - | - | 1 |
| **Or-I** | 44 | - | - | 44 |
| **Or-II** | 28 | - | - | 28 |
| **Or-III** | 16 | - | - | 16 |
| **YYT Landraces** | - | - | 92 | 92 |
| **Total** | **113** | **103** | **92** | **308** |

The resulting phylogenetic tree (Fig. SI1.1) showed that the YYT rice accessions consisted of two main groups, one closely related to the *indica* worldwide ecotype, the other one to the *japonica* worldwide ecotype, which was consistent to previous studies (Jiao et al, 2012; Liao et al. 2016). Specifically, the *indica* traditional landraces from YYT seemed to be related to, but genetically distinct from the worldwide *indica* gene pool. Further analyses including a larger set of YYT representatives and full genome data are needed to test this hypothesis.

**Figure SI1.1. Maximum-likelihood total evidence phylogenetic tree of 92 YYT accessions (black dots) and 213 worldwide accessions of *O. sativa* and related *Oryza* species (colored dots and squares) built from 44,855 SNPs using RAxML.**

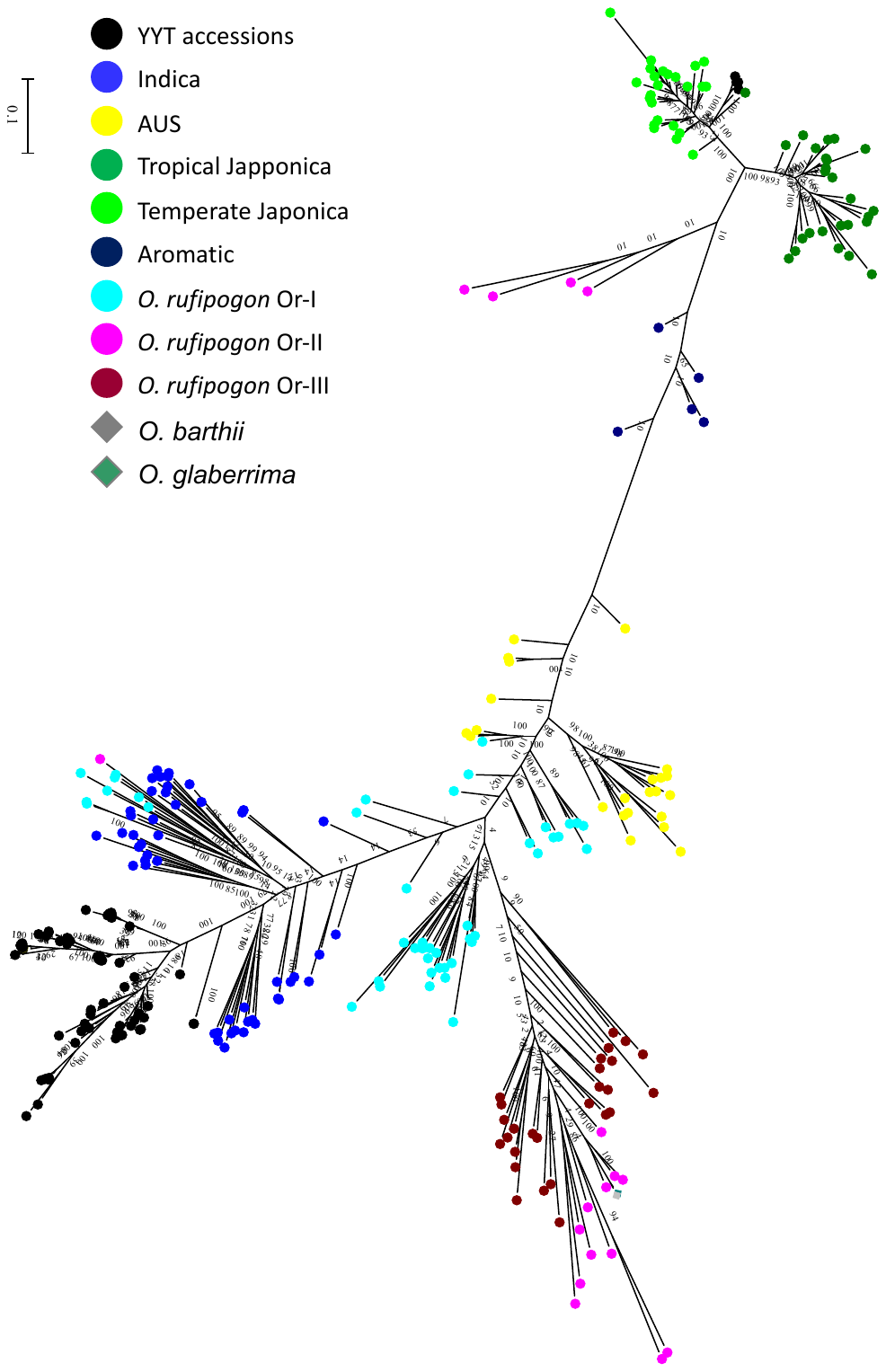
